# Transient non-specific DNA binding dominates the target search of bacterial DNA-binding proteins

**DOI:** 10.1101/2020.08.13.249771

**Authors:** Mathew Stracy, Jakob Schweizer, David J Sherratt, Achillefs N Kapanidis, Stephan Uphoff, Christian Lesterlin

## Abstract

Despite their diverse biochemical characteristics and functions, all DNA-binding proteins share the ability to accurately locate their target sites among the vast excess of non-target DNA. Towards identifying universal mechanisms of the target search, we used single-molecule tracking of 11 diverse DNA-binding proteins in living *Escherichia coli*. The mobility of these proteins during the target search was dictated by DNA interactions, rather than by their molecular weights. By generating cells devoid of all chromosomal DNA, we discovered that the nucleoid does not pose a physical barrier for protein diffusion, but significantly slows the motion of DNA-binding proteins through frequent short-lived DNA interactions. The representative DNA-binding proteins (irrespective of their size, concentration, or function) spend the majority (58-99%) of their search time bound to DNA and occupy as much as ∼30% of the chromosomal DNA at any time. Chromosome-crowding likely has important implications for the function of all DNA-binding proteins.

## INTRODUCTION

DNA is organized into chromosomes that must be maintained in a highly compacted state, while keeping the genetic information accessible for processing by many DNA-binding proteins. The ability of these proteins to identify and bind to specific DNA target sites among the vast excess of non-target DNA is crucial to fundamental cellular functions, including the recruitment of transcription factors to promoter sequences, of DNA repair proteins to DNA lesions, or of DNA topoisomerases to supercoiled DNA strands, to name just a few. In all organisms, diffusion is the primary mechanism by which DNA-binding proteins locate their target sites on chromosomes (Erbaş and Marko, 2019; Erbaş et al., 2019; Schavemaker et al., 2018). The diffusion coefficient of a particle in a dilute solution is determined by its size, as well as the viscosity and temperature of the medium. Within the crowded and heterogeneous intracellular environment, however, a myriad of specific and non-specific interactions as well as steric effects influence the mobility of macromolecules. Because of this complexity, efforts to understand molecular mobility have relied on phenomenological models (Kalwarczyk et al., 2012; Mika and Poolman, 2011) or coarse-grained simulations of the cytoplasm (Chow and Skolnick, 2017; Feig et al., 2015; Hasnain et al., 2014). In this context, analysis of *in vivo* experimental data is crucial not only to determine parameter values but also the structure of such models by informing which cellular components and interactions should be included in a model.

Contrary to eukaryotes, bacterial chromosomes are not compartmentalized into a nucleus, but organized into nucleoid structures without a physical barrier from the cytoplasm. The 4.6 Mbp *E. coli* chromosome with a contour length of 1.6 mm is compacted into a volume of ∼1 µm^3^ via DNA supercoiling, entropic forces, as well as protein-DNA and RNA-DNA interactions, and occupies ∼60% of the bacterial cell volume (Gray et al., 2019). Outside the nucleoid, the cytoplasm is mainly comprised of RNA and proteins. A longstanding question is whether the presence of the dense nucleoid mesh affects the mobility of all cytoplasmic proteins, regardless of their ability to bind DNA. The chromosome could pose a steric barrier, resulting in confined diffusion, and preventing larger proteins from accessing the densest regions of the nucleoid (Kalwarczyk et al., 2012; Konopka et al., 2006; Kuznetsova et al., 2014). Furthermore, the target-search process is subject to a trade-off between speed and accuracy to distinguish target from non-target sites (Zandarashvili et al., 2015). Accumulating experimental evidence supports theoretical considerations that the search efficiency is maximized by “facilitated diffusion”, which is the combination of 3D protein diffusion with non-specific binding and 1D sliding along DNA (Halford and Marko, 2004; Hammar et al., 2012; Hippel and Berg, 1989). Together with chromosome crowding effects, the relative contribution of 3D and 1D diffusion modes during the target search should strongly affect the overall mobility of DNA-binding proteins *in vivo*.

Fluorescence microscopy-based methods such as Fluorescence Correlation Spectroscopy (FCS) (Bacia et al., 2006; Cluzel et al., 2000) and Fluorescence Recovery After Photobleaching (FRAP) (Konopka et al., 2006; Kumar et al., 2010; Mika and Poolman, 2011; Mika et al., 2010; Mullineaux et al., 2006; Nenninger et al., 2010; Ramadurai et al., 2009) have been used to investigate protein mobility in live bacterial cells. More recently, it has become possible to directly visualize aspects of the target search of individual proteins in live cells using single-molecule microscopy (Elf et al., 2007; Hammar et al., 2012; Kapanidis et al., 2018; Normanno et al., 2015; Rhodes et al., 2017). These studies focused on a limited number of test proteins, typically the Lac repressor or other transcription factors, raising the question whether the proposed models for the target search are universal for diverse types of DNA-binding proteins. Although the observed intracellular mobility and spatial distribution of DNA-binding proteins suggest that non-specific DNA interactions play an important role in their target search kinetics, these interactions appeared too transient for direct visualization and quantification by live-cell imaging (Garza de Leon et al., 2017; Stracy et al., 2015, 2016; Uphoff et al., 2013). In previous attempts to resolve this issue, the DNA-binding affinity of the protein studied was perturbed genetically (Elf et al., 2007), but for some proteins this approach is not readily tractable. Alternatively, protein mobility has been compared between different regions of the cell with lower or higher DNA density (Bakshi et al., 2011; Sanamrad et al., 2014; Stracy et al., 2015). However, since few DNA-binding proteins are located in DNA-free regions of the cell, it is difficult to accurately measure their diffusion with this approach. Furthermore, even with super-resolution microscopy, the exact shape and boundary of the nucleoid are not well defined (Le Gall et al., 2017; Stracy et al., 2015).

To overcome this uncertainty and to determine the influence of the nucleoid DNA on protein mobility, we measured the mobility of DNA-binding proteins in cells devoid of chromosomal DNA. To this end, we developed a method to remove all chromosomal DNA from cells. By comparing protein diffusion in DNA-free cells and unperturbed cells, we identified universal features of the target search process for 11 DNA-binding proteins with a broad range of sizes, biochemical characteristics and functions. This, combined with diffusion simulations, allowed us to quantitatively partition the behavior of diverse DNA-binding proteins into long-lived DNA-binding at target sites, transient non-specific DNA-binding, and free diffusion between DNA strands. We found that the intracellular mobility of proteins during their target search is primarily dictated by transient interactions with the DNA, rather than by their molecular weight or intracellular concentration. The representative DNA-binding proteins (irrespective of their size, concentration, or function) spend the majority (58-99%) of their search time bound to DNA, occupying as much as ∼30% of the chromosomal DNA at any time.

## RESULTS

### Live-cell single-molecule tracking of a variety of DNA-binding proteins

To uncover universal mechanisms that govern the target search process of DNA-binding proteins in general, we measured the diffusion characteristics of 11 different proteins involved in various types of DNA transactions and spanning a large range of molecular weights and concentrations inside the cell. These included proteins whose target is a specific DNA sequence such as RNA polymerase (β’ subunit, RpoC), the low copy number transcription factor LacI, and the abundant histone-like nucleoid-associated proteins HU and H-NS. We further analyzed proteins which target DNA structural motifs such as DNA topoisomerases (ParC, GyrA) which act on supercoiled DNA, or DNA polymerase I (Pol1) and DNA ligase (LigA) which recognize gapped or nicked DNA respectively. Lastly, we also studied DNA-repair proteins which recognize DNA lesions (UvrA) or mismatches (MutS), and the Structural Maintenance of Chromosomes (SMC) protein MukB which is involved in chromosome organization but binds DNA with little known specificity.

To examine the mobility of this diverse set of proteins, we use single-molecule tracking, a method that provides a direct readout of protein mobility inside living cells (Gahlmann and Moerner, 2014; Li et al., 2018; Uphoff and Sherratt, 2017). We imaged proteins that were fused to the photoactivatable fluorescent protein PAmCherry (Subach et al., 2009), and expressed from their endogenous chromosome locus in *Escherichia coli* cells. The use of a photoactivatable fluorophore allows tracking proteins at their native expression levels by imaging single molecules, one at a time, while the rest of the molecules reside in a non-fluorescent state (Bakshi et al., 2011; English et al., 2011; Manley et al., 2008; Niu and Yu, 2008; Uphoff et al., 2013). We recorded movies on a custom-built microscope using near-Total Internal Reflection Illumination (Tokunaga et al., 2008; Wegel et al., 2016) with sparse photoactivation at a frame rate of 15 ms/frame for >5000 frames to resolve the motion of hundreds of molecules per cell in multiple cells per field of view. Following automated localization and particle tracking analysis, the apparent diffusion coefficient D* was calculated from the mean-squared displacement (MSD) of each trajectory (Uphoff, 2016) (Fig. 1A).

**Figure 1.**
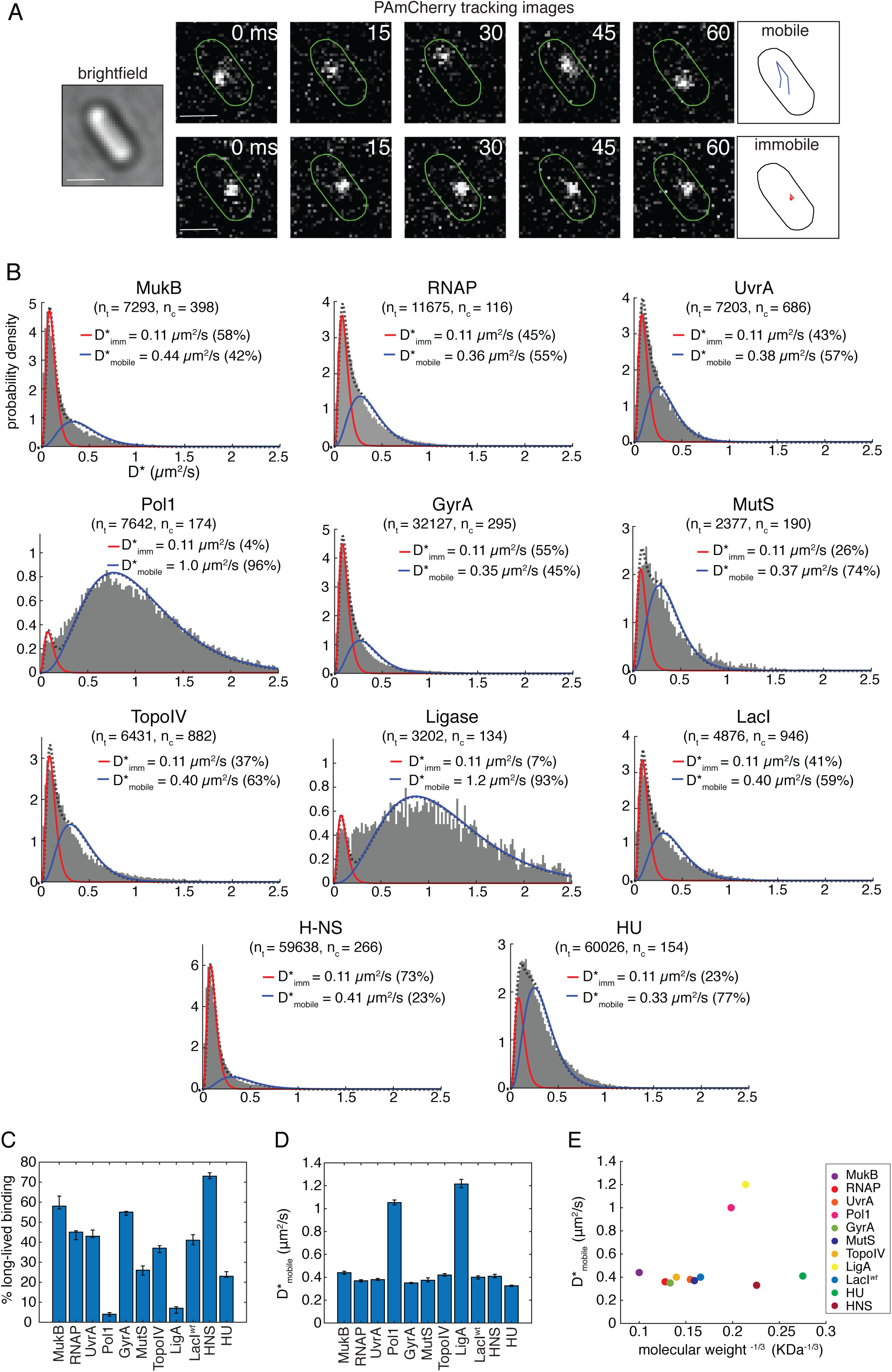
Intracellular mobility of diverse types of DNA-binding proteins in live *E. coli* cells is highly variable and unrelated to their molecular weights. (A) Illustration of photoactivated single-molecule tracking, showing example fluorescence images and trajectories of a mobile molecule (top) and immobile molecule (bottom) within the cell (green outline). Scale bar 1 µm. (B) Histograms of apparent diffusion coefficients D* (grey bars) for diverse DNA-binding proteins, fitted with a model (black dashed line) of a mixture of immobile (red) and mobile molecules (blue). The number of cells (n_c_ =) and the numbers of tracks (n_t_ =) analyzed are indicated. (C) Percentages of long-lived binding molecules obtained from fitting D* histograms in Fig. 1 with a model of a mixture of immobile and mobile molecules. Error bars represent 95% confidence intervals. (D) D_mobile_* values for the mobile molecule populations. (E) D_mobile_* plotted against the cubic root of the molecular weight of the protein complex.

Computing D* from particle-tracking data tends to underestimate the actual diffusion coefficient D for molecules with high mobility because of measurement biases, including the blurring of fluorescent spots due to molecular motion during the camera exposure and the confinement of trajectories within the cell volume (Uphoff, 2016). Hence, we initially analyzed apparent diffusion coefficients and subsequently employed diffusion simulations to correct the biases (see below). Molecular subpopulations that differ in mobility can be detected as separate species in D* distributions. In particular, proteins that remain bound to DNA over the entire trajectory appear essentially immobile due to the slow and constrained motion of chromosomal DNA (Elmore et al., 2005).

### The mobility of DNA-binding proteins is independent to their molecular weights

The average intracellular mobility of the different DNA-binding proteins varied strongly, ranging from mostly immobile proteins such as HNS (mean D* = 0.17 µm^2^/s) to mostly diffusing proteins such as LigA (mean D* = 1.14 µm^2^/s) (Fig. 1B). There was no obvious relation between the observed mobility and the type of DNA interactions (e.g. sequence-specific, structure-specific, or lesion binding). To distinguish between proteins specifically bound to DNA and mobile proteins searching for target sites we first determined the apparent mobility of proteins specifically bound to DNA by measuring the motion of Pol1 molecules recruited to DNA damage sites generated by treating cells with the DNA-alkylating agent methyl methanesulfonate (MMS, 100 mM) (Fig. S1A), which resulted in an increase in molecules which are immobile (with D_imm_*=0.11 µm^2^/s) for the entire trajectory (5 frames, 75 ms), as previously observed (Uphoff et al., 2013). For other proteins in this study we have previously observed a similar increase in ‘long-lived’ immobile molecules (D_imm_*=0.11 µm^2^/s) upon recruitment to specific target sites after induction of DNA damage (LigA, UvrA, MutS; Stracy et al., 2016; Uphoff et al., 2013, 2016), or by capturing DNA-bound enzymes during catalysis by drug treatment (GyrA, ParC; Stracy et al., 2019; Zawadzki et al., 2015). We previously observed a decrease in the long-lived immobile population of RNAP upon addition of a transcription-inhibiting drug (Stracy et al., 2015), and a similar decrease for LacI after removing its chromosomal binding site (Garza de Leon et al., 2017). Together, these studies show that the long-lived immobile population represents proteins specifically bound at DNA target sites. The apparent mobility of these DNA-bound molecules was slightly above the localization uncertainty of σ= 35 nm measured in chemically fixed cells (giving an apparent D_fixed_*=0.07 µm^2^/s; Fig. S1B). By subtracting the contribution of the localization uncertainty to the observed D* (Michalet and Berglund, 2012), we estimate the mobility of proteins bound to DNA loci for the entire trajectory as D*_bound_ = 0.04 µm^2^/s.

To determine the relative abundances and average diffusion coefficients of mobile molecules searching for target sites and long-lived immobile molecules bound to DNA, we fitted the D* histograms using an analytical function derived from a two-species Brownian motion model (Stracy et al., 2015) (Fig. 1B and S1A-B). The quantification confirmed our initial observations that the different DNA-binding proteins exhibit vastly different mobility inside cells, both in terms of the fraction of mobile and immobile molecules (ranging from 96 % of mobile for Pol1 to 23 % mobile for H-NS molecules) (Fig. 1C), and in terms of the diffusion coefficients of the mobile molecules (ranging from 0.33 µm^2^/s for HU to 1.2 µm^2^/s for LigA) (Fig. 1D).

According to the Stokes-Einstein equation for Brownian motion, the diffusion coefficient of a spherical particle is related to its mass: D ∼ M^-1/3^. To test this relation for the DNA-binding proteins, we plotted D* of molecules in the mobile state against the known molecular weights of each protein (Fig. 1E). Strikingly, the *in vivo* mobility of DNA-binding proteins was largely independent of their mass. Although non-spherical proteins are expected to deviate from the Stokes-Einstein law, this does not explain the absence of any correlation between mass and mobility. In contrast, previous studies showed a clear dependence of mass on the mobility of cytoplasmic proteins with no affinity for DNA (Kalwarczyk et al., 2012; Kumar et al., 2010; Nenninger et al., 2010). Our results indicate that the apparent mobility of DNA-binding proteins is dictated by molecular interactions independent of protein mass. There was also no trend between the mobility and the intracellular concentration of the different proteins (Fig. S1C).

### DNA-binding proteins remain closely associated with the nucleoid during their target search

We examined the spatial distribution of mobile DNA-binding proteins relative to the nucleoid. As an example, we tracked Pol1-PAmCherry and RNAP-PAmCherry in live cells that were stained with SytoGreen dye to label DNA (Fig. 2A-B). The spatial distributions of mobile molecules closely overlapped with the nucleoid. Similarly, when averaged over many cells and the intracellular position of the mobile population of molecules clearly demarcates the nucleoid shape (Fig. 2D). This was in contrast to ribosomal protein S1, which has no direct DNA affinity. Consistent with previous reports (Sanamrad et al., 2014), the slow-moving S1 molecules, which are presumably incorporated into ribosomes, resided outside the nucleoid area, whereas the mobile unincorporated subunits are uniformly distributed throughout the cell (Fig. 2C-D). We hypothesized that the enrichment of mobile DNA-binding proteins within the nucleoid is caused by transient interactions with DNA during the target search process. The computation of D* values is based on the average movement of a molecule over a series of frames (here 5 frames, 75 ms). The observed mobility thus reflects a time-average of the diffusion coefficient where 3D diffusion is interrupted by multiple transient binding events with a duration below 75 ms. Consistent with this view, we observed that the mobility of DNA-binding proteins increased and tracks spread throughout the cell cytoplasm after treatment with the antibiotic rifampicin, which causes decompaction of the nucleoid (Fig. S1D) (Cabrera et al., 2009; Dworsky and Schaechter, 1973; Stracy et al., 2015).

**Figure 2.**
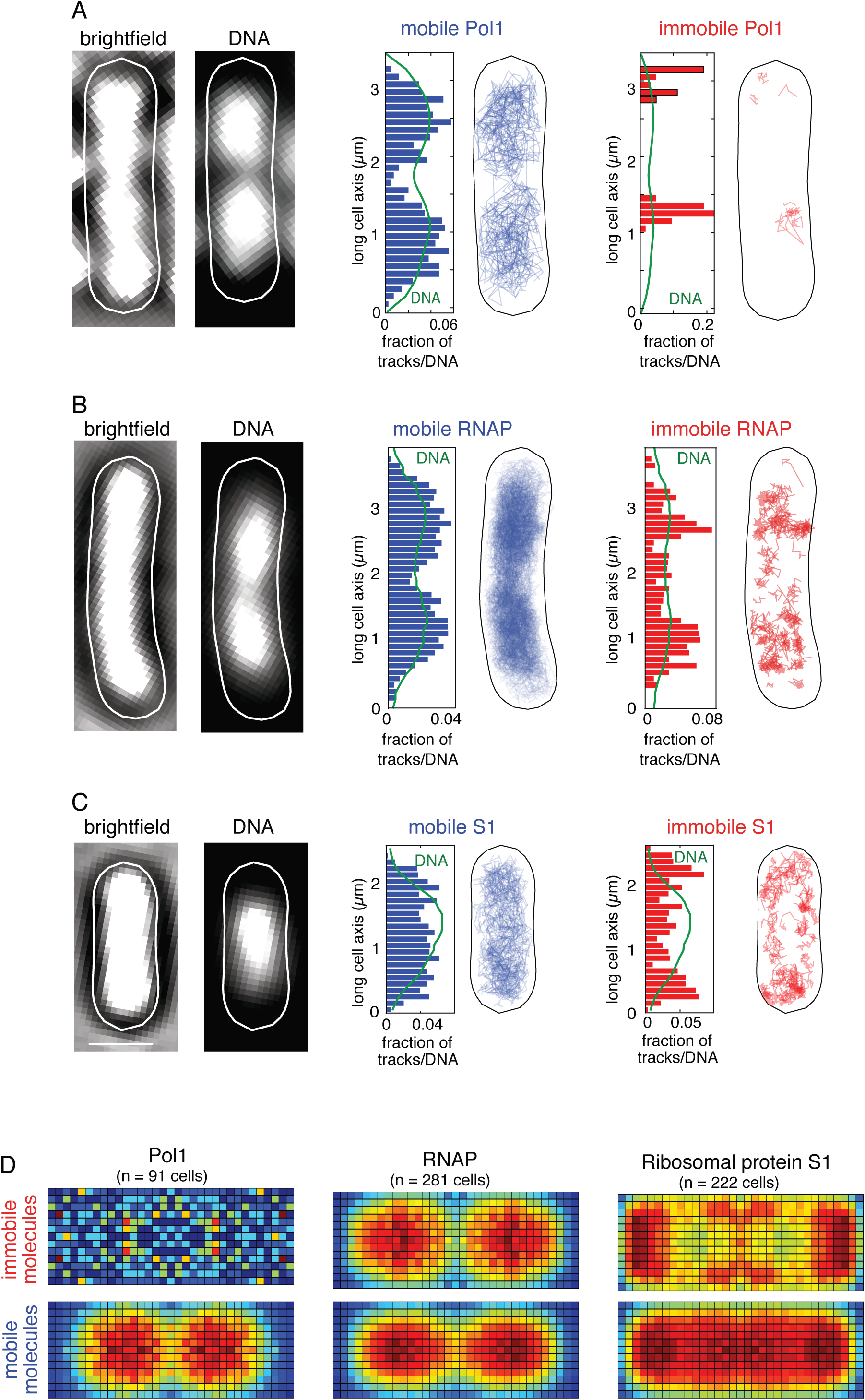
DNA-binding proteins stay closely associated with the nucleoid during the target search. (A) Localizations of Pol1 (PolA-PAmCherry), (B) RNAP (RpoC-PAmCherry), and (C) ribosomal protein S1 (S1-PAmCherry) molecules relative to the nucleoid. From left to right: transmitted light image (scale bar 1 µm); SytoGreen-stained nucleoid DNA with segmented cell outline; maps showing the tracks of mobile (blue) and immobile (red) PAmCherry fusion proteins in cells. Histograms show localizations across the long cell axis (blue/red bars) together with the SytoGreen fluorescence profile of the nucleoid (green line). (B) Average spatial distributions of Pol1, RNAP and ribosomal protein S1 immobile and mobile molecules.

### Generating chromosome-free cells to study protein diffusion in the absence of DNA

The association of mobile DNA-binding proteins with the nucleoid could reflect a genuine DNA-binding activity or be the result of a sieving effect where protein movement is slowed within the nucleoid by physical entrapment within the mesh of DNA strands. To distinguish between the effects of sieving and non-specific DNA interactions, we devised a method that eliminates protein-DNA interactions entirely by removing all chromosomal DNA from cells, while retaining the same cell size and intracellular protein concentration. To this end, we used the I-SceI restriction endonuclease from *Saccharomyces cerevisiae* (Monteilhet et al., 1990), which introduces site-specific double-stranded DNA breaks (DSBs) at *I-SceI* cut sites (*I-SceI*^*cs*^) inserted into the *E. coli* chromosome (Fig. 3A) (Lesterlin et al., 2013; Meddows et al., 2004). In the absence of RecA, which is essential for homologous recombination, creation of DSBs by I-SceI results in complete degradation of the chromosome by the RecBCD helicase-nuclease complex, a phenomenon referred to as *reckless* chromosome degradation (Skarstad and Boye, 1993; Willetts and Clark, 1969). To minimize the time required for complete chromosome degradation, we inserted two cut sites diametrically opposed on the genetic map of the chromosome: in the *ilvA* locus (3953 kb) close to the origin of replication, and in the *ydeO* locus (1580 kb) in the *terminus* region (referred to as OT strain) (Fig. 3A). We then inactivated the *recA* gene by mutation in these strains carrying *Origin*-*Terminus* cut sites (referred to as OT*recA-*). Chromosome degradation was triggered by the expression of the plasmid-borne *I-SceI* gene under the control of an arabinose-inducible promoter. Chromosome degradation after *I-SceI* induction resulted in the progressive disappearance of DAPI-stained DNA from cells (Fig. 3B), which was complete within 120 to 160 min in most (∼92%) cells (Fig. 3C, Fig. S2A-B). This reflects the time required for 4 RecBCD complexes to each degrade approximately one quarter of the chromosome (∼1150 kb) from the 4 DNA ends generated by 2 DSBs at a speed of ∼160 bp per second, consistent with previous results (Lesterlin et al., 2014). Before arabinose induction, a fraction of cells (∼17%) already exhibited DNA degradation due to leaky I-SceI expression (Fig. S2A-B). We also found that a fraction (∼8 %) of cells did not exhibit complete chromosome loss after 120 min (Fig. S2A-B), likely due to heterogeneous induction of I-SceI from the arabinose-inducible promoter in the cell population (Siegele and Hu, 1997) or because of the limiting number of RecBCD molecules per cell (Lepore et al., 2019). In the following, to ensure our results reflected completely chromosome-free cells, we excluded cells that showed any remaining fluorescent DNA stain from our analysis.

**Figure 3.**
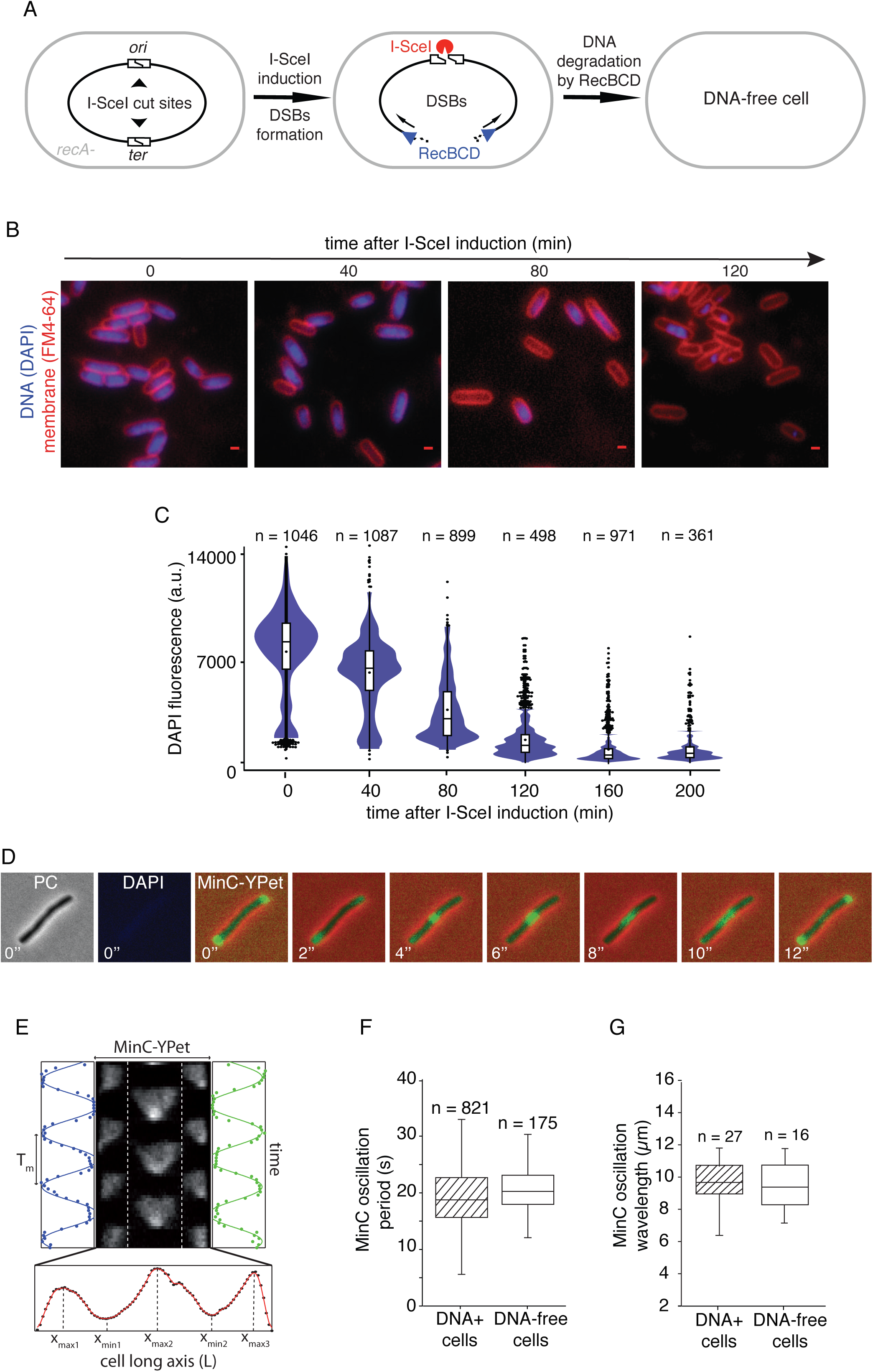
Generating chromosome-free cells that remain metabolically active. (A) Schematic of the chromosome-degradation system. The production of I-SceI endonuclease induces the formation of two double-stranded-breaks (DSBs) at *I-SceI* cut-sites inserted at diametrically opposed position on the chromosome. In a *recA-* mutant strain, processing of the two DSBs by RecBCD complexes results in the complete degradation of the chromosome. (B) Chromosome degradation following I-SceI induction is revealed by the loss of DAPI-stained DNA fluorescence (blue) in cells with FM464-labelled membrane (red). Scale bar, 1 µm. (C) Distributions of DAPI fluorescence shows complete chromosome degradation 120 min after I-SceI induction (black dot: mean and outliers, horizontal lines: median, 1^st^ and 3^rd^ quartiles). The number of cells analyzed (n) is indicated. (D) MinC-YPet oscillation in an example chromosome-free cell. Cell filamentation was induced by cephalexin treatment. Transmitted light, DAPI, and MinC-YPet fluorescence images (2 s/frame time lapse) obtained 120 min post I-SceI induction. Scale bar, 1 µm. (E) Kymograph of MinC-YPet oscillation in an example cephalexin-treated filamentous cell. The width of the kymograph corresponds to the cell long-axis (L). Time-dependent intensity signals in the cell halves are shown in blue and green with the oscillation period T_m_. Time-average concentration profile shown underneath for measuring the oscillation wavelength. (F, G) MinC-YPet oscillation period and wavelength are similar in filamentous cells with and without chromosome degradation. The number of cells analyzed (n) is indicated.

### Chromosome-free cells remain metabolically active for several hours

Since protein mobility is influenced by the metabolic state of the cell (Parry et al., 2014), we explored if cells remained metabolically active after chromosome degradation using two independent assays. First, to test if ATP-driven mechanisms were affected by chromosome loss, we turned to the well-characterized ATP-dependent Min system, whose three components MinCDE are important for defining the position of the division site (Lutkenhaus, 2007). Coupled protein–protein and protein– membrane interactions generate pole-to-pole dynamic oscillation of MinC, which is highly sensitive to ATP concentrations (Hu et al., 2002). These oscillations are particularly striking in cells that have been grown into long filaments by treatment with the antibiotic cephalexin (Raskin and de Boer, 1999). We found that the MinC-Ypet oscillation period was ∼17 seconds with a wavelength of ∼10 µm, both in unperturbed cells and after chromosome degradation (Fig. 3D-G and Fig. S3), demonstrating that ATP concentration in chromosome-free cells remained stable for at least 2 hours. Our results also indicate that the presence of the nucleoid DNA has no influence on Min protein dynamics, in contrast to a previous report that the oscillations may be coupled to chromosome segregation (Di Ventura et al., 2013).

Second, to test if protein synthesis activity was maintained after chromosome loss, we used a non-degraded plasmid producing a reporter protein ParB-mCherry from a *P*_*lac*_ promoter. We found that IPTG-induced ParB-mCherry production continued for ∼200 min after I-SceI induction (Fig. S4A). Together, these tests establish that our chromosome degradation strategy is appropriate to study protein diffusion in metabolically active chromosome-free cells. Furthermore, in order to validate the use of this genetic system, we confirmed that protein diffusion was not affected by inactivation of RecA *per se*, or by induction of I-SceI in cells that do not contain any I-SceI cut sites, nor by DSB creation in RecA+ DNA repair-proficient cells (Fig S4B-C).

### The mobility of the Lac repressor increases in DNA-free cells

To test the effect of chromosome loss on intracellular diffusion, we first focused on the lac repressor (LacI) as a prototypical DNA-binding protein which searches for operator sequences by facilitated diffusion involving frequent non-specific DNA binding, rotation-coupled sliding and hopping (Elf et al., 2007; Garza de Leon et al., 2017; Hammar et al., 2012; Kao-Huang et al., 1977; Marklund et al., 2020). Chromosome degradation, ∼120 min after I-SceI induction, drastically changed the diffusion behavior of LacI-PAmCherry (Fig. 4A). We no longer detected any immobile molecules; further, the mobility of the diffusing population increased significantly (from D* = 0.43 µm^2^/s in unperturbed cells to D* = 1.5 µm^2^/s in chromosome-free cells). The change in diffusion pattern is apparent in the D* distribution (Fig. 4A), MSD curves (Fig. 4B), and cumulative distributions of displacements (Fig. 4C). We identified a subpopulation of cells (∼15 %) that exhibited no change in LacI diffusion after I-SceI induction; however, DNA staining prior to single-molecule tracking showed that these cells had no or incomplete chromosome loss and these cells were thus excluded (Fig. S5A-B). The strong influence of the presence of the chromosome on LacI mobility could be due to DNA-binding and sliding, or the result of a general molecular sieving effect, where protein motion is hindered because of entrapment within the chromosome meshwork. The latter effect should influence the motion of all proteins in the cell, even those that have no DNA affinity. To test this directly, we imaged a truncated LacI^-41^ mutant with most of its DNA-binding domain (41 amino acids from the N-terminus) removed. For this mutant all specific and non-specific DNA binding modes are abolished (Elf et al., 2007; Garza de Leon et al., 2017), and hence shows essentially no immobile molecules (Fig. 4D). Notably, LacI^-41^ also had a much higher apparent diffusion coefficient than the mobile population of wild-type LacI (D_lacI-41_*=1.3 µm^2^/s vs D_lacI_*=0.43 µm^2^/s). This difference far exceeded the 2–3% change expected solely from the 9 kDa decrease in the protein size due to the truncation (considering D ∼ M^-1/3^). After chromosome degradation, LacI^-41^ only showed a small increase in mobility (from D*=1.3 µm^2^/s to 1.5 µm^2^/s) (Fig. 4B-D). To test if this is general, we also measured the diffusion of unconjugated PAmCherry alone and found no significant change between unperturbed and chromosome-free cells (Fig. 4C). Therefore, the presence of the chromosome has only a minor influence on the diffusion of a protein that has no affinity for DNA. This is consistent with observations that the fluorescent proteins mEOS and Kaede diffuse inside the whole cell volume with no evidence that the presence of the nucleoid hinders their motion (Bakshi et al., 2011; English et al., 2011). Note that these data do not exclude the possibility that DNA sieving may hinder the movement of proteins and macromolecular complexes that are much larger than LacI and fluorescent proteins, such as 70S ribosomes which are occluded from the nucleoid (Fig. 2C, and Sanamrad et al., 2014).

**Figure 4.**
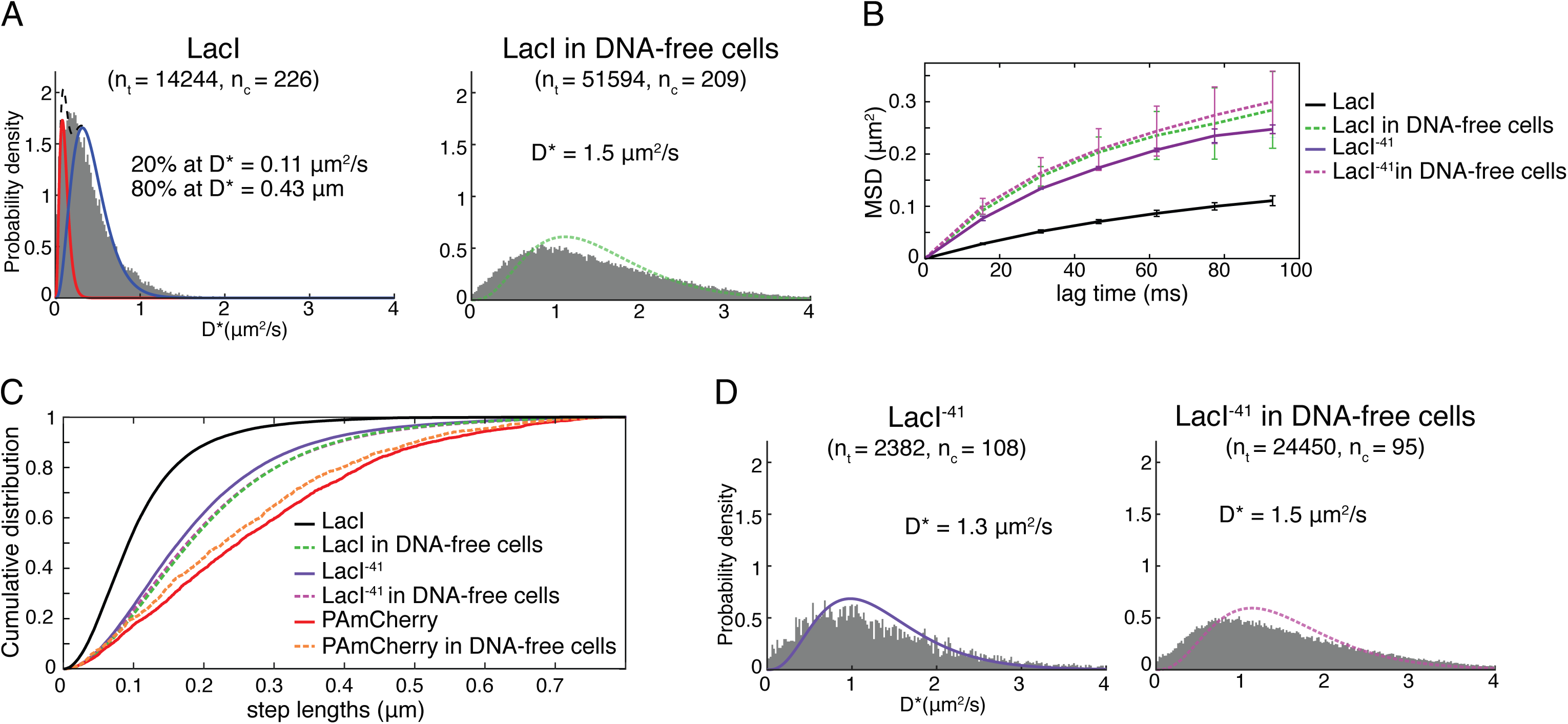
Diffusion of lac repressor increases in chromosome-free cells. (A) D* histograms of LacI-PAmCherry in unperturbed cells (left) fitted with a model (black dashed line) of a mixture of immobile (red) and mobile molecules (blue). D* distribution of LacI-PAmCherry in chromosome-free cells 120 min after I-SceI induction (right) fitted with a model for mobile molecules (green).). (B) Mean squared displacement plots corresponding to data in panels (A) and (D). (C) Cumulative distributions of the step lengths between consecutive localizations in unperturbed and in chromosome-free cells for LacI-PAmCherry, LacI^41^-PAmCherry, and unconjugated PAmCherry. The distributions shift to longer steps with increasing diffusion coefficient. (D) D* histograms of mutant LacI^41^-PAmCherry in unperturbed cells (left) and in chromosome-free cells 120 min after I-SceI induction (right) fitted with a model for mobile molecules (purple and magenta, respectively.

### Transient DNA interactions strongly affect the mobility of diverse DNA-binding proteins

Having established that chromosome degradation increases the mobility of LacI primarily because of a loss of DNA interactions, we asked if this was generally the case for diverse types of DNA-binding proteins. We chose four proteins representing distinct types of DNA interactions (Fig. 5A): RNA polymerase (RNAP) recognizes specific promoter sequences to initiate RNA synthesis and transcribe genes (Mazumder and Kapanidis JMB 2019); DNA polymerase I (Pol1) recognizes gapped or nicked structures in DNA repair and replication (Joyce and Steitz, 1994); Structural Maintenance of Chromosomes protein MukB interacts non-specifically with double-stranded DNA to aid chromosome segregation (Nolivos et al., 2016; Reyes-Lamothe et al., 2012; Rybenkov et al., 2014); Ligase (LigA) interacts with DNA nicks and catalyzes the joining of DNA ends (Shuman, 2009). These proteins not only have different biological functions, but also differ in their shapes, molecular weights, oligomeric states, and intracellular concentrations. The RNAP holoenzyme with the initiation factor σ^70^ is a 449 kDa complex composed of 6 different proteins and present at 3000-6000 copies per cell (Bakshi et al., 2013; Endesfelder et al., 2013; Stracy et al., 2015). Pol1 is a monomeric 104 kDa protein with two globular domains connected by a flexible linker and present at ∼500 copies per cell (Uphoff et al., 2013). MukB has a characteristic elongated SMC protein fold with globular domains on either end of 50-nm long coiled-coil domains. Approximately 100 MukB homodimers per cell likely form large 890 kDa complexes with MukE and MukF proteins (Badrinarayanan et al., 2012). Ligase is a monomeric 73 kDa enzyme present at a ∼100 copies per cell that encircles the DNA as a C-shaped protein clamp (Nandakumar et al., 2007; Shuman, 2009).

**Figure 5.**
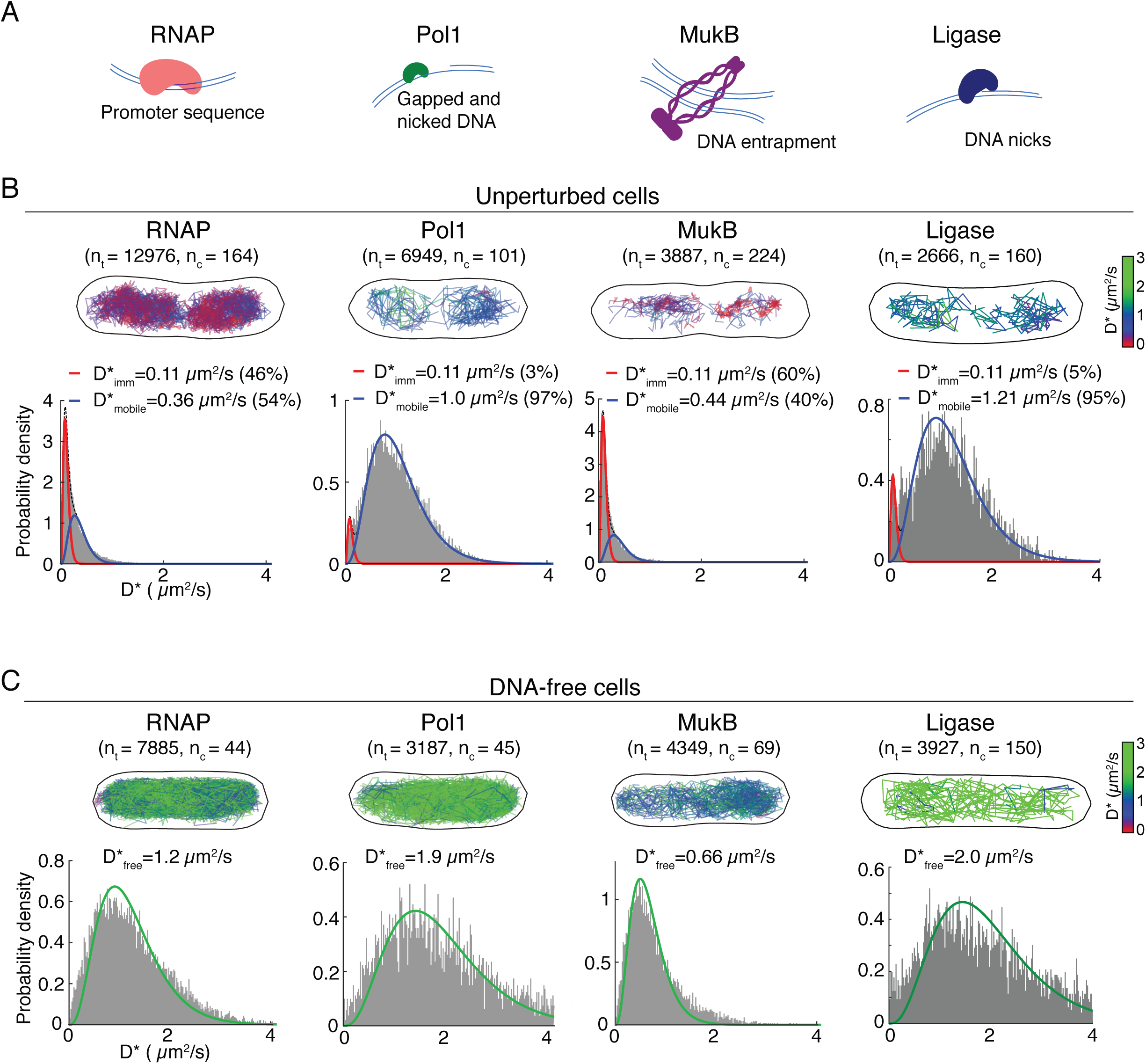
Chromosome degradation increases the mobility of diverse types of DNA-binding proteins. (A) Diagrams of RNAP, Pol1, MukB and Ligase DNA-binding properties. (B) Tracks of RNAP-PAmCherry, Pol1-PAmCherry, MukB-PAmCherry and LigA-PAmCherry in example cells, with the color of each track representing its D* value. D* histograms in cells without I-SceI induction, fitted with a model of a mixture of immobile (red) and mobile (blue) molecules. (B) Tracks of RNAP-PAmCherry, Pol1-PAmCherry, MukB-PAmCherry and LigA-PAmCherry in example chromosome-free cells 120 min after I-SceI induction with the color of each track representing its D* value. D* histograms in chromosome-free cells fitted with a model for mobile molecules. The number of cells (n_c_=) and the numbers of tracks (n_t_ =) analyzed are indicated in B and C.

Considering the differences in the function and physical characteristics of RNAP, Pol1, MukB, and LigA, any shared aspects of their diffusion behavior are likely to indicate universal mechanisms of the DNA target search. In unperturbed cells, a large fraction of RNAP-PAmCherry and MukB-PamCherry molecules were immobile or slowly diffusing (RNAP: D* = 0.36 µm^2^/s, MukB: D* = 0.39 µm^2^/s), whereas Pol1-PAmCherry and LigA-PAmCherry molecules were rarely immobile for the entire trajectory and diffused faster (D* = 1.0 µm^2^/s and D* = 1.2 µm^2^/s respectively) (Fig. 5B), consistent with our previous observations (Badrinarayanan et al., 2012; Stracy et al., 2015; Uphoff et al., 2013). Despite the differences in the diffusion profiles, a unifying feature was the clear nucleoid-association of the tracks for all three proteins in unperturbed cells (Fig. 5B; Fig. 2). Chromosome degradation had the same effects for all four proteins (compare Fig. 5B and C): the populations of long-lived immobile molecules disappeared, and diffusion of the mobile proteins increased substantially (RNAP: D* = 1.2 µm^2^/s, Pol1: D* = 1.9 µm^2^/s, MukB: D* = 0.66 µm^2^/s, MukB: D* = 1.9 µm^2^/s) (Fig. 5C). Furthermore, the tracks filled the entire cytoplasm of chromosome-free cells (Fig. 5C). These results match our observations for the Lac repressor (Fig. 4), and taken together, they demonstrate that transient DNA interactions dictate the mobility and spatial distribution of diverse types of DNA-binding proteins.

### The mobility of DNA-binding proteins shows a steep size-dependence in chromosome-free cells

Accurate quantification of diffusion coefficients from single-molecule tracking experiments requires consideration of several biases such as localization error and confinement within the cell volume (especially for rapidly moving molecules) (Uphoff, 2016). In order to account for these potential biases to determine accurate *D* values from experimentally measured *D*\*, we applied stochastic Brownian motion simulations to generate artificial single-molecule tracks using an identical number of molecules inside the same segmented 3D cell volumes as in the experimental data (Fig. 6A). Localization error and stochastic disappearance of tracks due to photobleaching were also modeled, resulting in the same sampling and biases as in the experimental *D*\* distributions. We determined an unbiased estimate of the diffusion coefficient *D* from the best match (according to a least-squares metric) between D* distributions observed in experiments and those obtained from simulations with a range of input diffusion coefficients.

**Figure 6.**
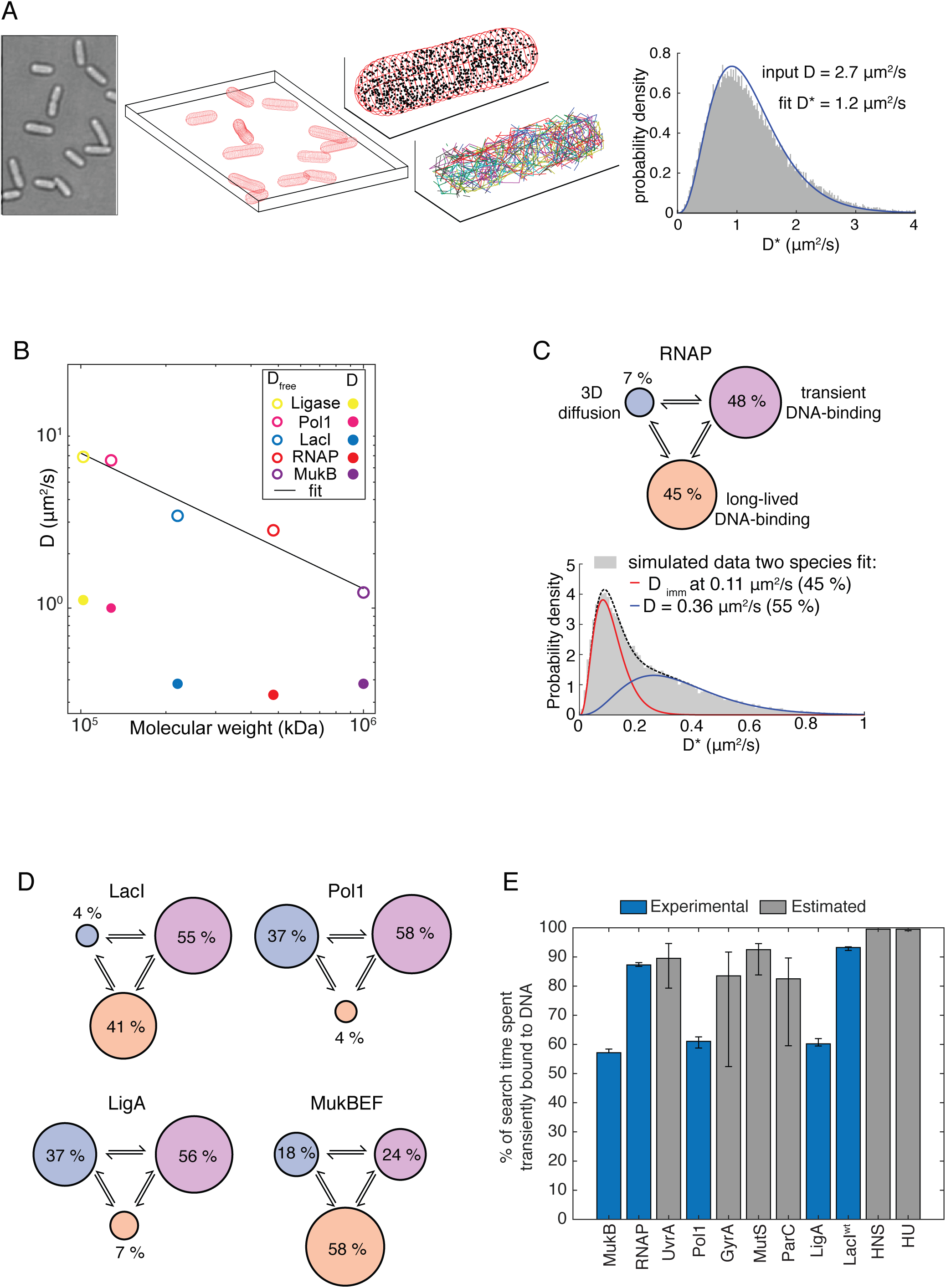
Quantitative partitioning of protein states. (A) Illustration of Brownian motion simulation to estimate unbiased diffusion coefficients D_mobile_ (in unperturbed cells) and D_free_ (in chromosome-free cells). (B) D_mobile_ and D_free_ plotted *vs* molecular weight M on log-scale. Linear fit log(D_free_) = α·log(c·M). (C) Partitioning long-lived DNA-binding (orange), transient DNA-binding (purple), and 3D diffusion (blue) states for RNAP and (D) for LacI, Pol1, LigA and MukB. (E) The percentage of search time spent nonspecifically bound to DNA for all 11 DNA-binding proteins studied. Blue bars show the proteins with *D*_*free*_ measured in chromosome-free cells, grey bars show proteins with *D*_*free*_ estimated from the fit in panel B. Error bars correspond to standard deviations.

Using this procedure, we estimated the mean unbiased diffusion coefficients of LacI, RNAP, Pol1, LigA, and MukB molecules after chromosome degradation (*D*_*free*_ values in Table 1). To verify that these values are robust with regards to the data acquisition and simulation parameters, we also performed single-molecule tracking experiments at three-fold shorter camera exposure times (5 ms) and obtained the same results from the corresponding simulations (Fig. S6). Although there was no correlation between the mass and the diffusion coefficient of DNA-binding proteins in unperturbed cells (*D*_*mobile*_), we found a clear inverse relation in chromosome-free cells (*D*_*free*_) (Fig. 6B). Fitting a power-law D_free_ = c·M^α^ yielded an exponent of α = -0.75, showing that protein mobility decreases more steeply with increasing mass than predicted by the Stokes-Einstein model (α = -0.33). Therefore, the crowded cytoplasm has sieving properties even in the absence of the chromosome meshwork. Indeed, the diffusion of DNA-binding proteins in DNA-free cells shows a similar mass dependence as cytoplasmic proteins that have no DNA-binding function in unperturbed cells (α = -0.7) (Mika and Poolman, 2011).

**Table 1.** Quantitative partitioning of DNA-binding protein activity.

| Protein | Function | Size of PAmCherry labelled protein or protein complex (kDa)<sup>1</sup> | Molecules/cell | $D_{mobile}^*$ ($\mu\text{m}^2/\text{s}$) | $D_{free}$ ($\mu\text{m}^2/\text{s}$) | % of long-lived DNA-bound molecules | % transient DNA-bound molecules | % freely diffusing molecules | Copy number reference |
| --- | --- | --- | --- | --- | --- | --- | --- | --- | --- |
| RNA Polymerase ($\beta'$ subunit) | transcription | 480 (holoenzyme) | 4,000 | $0.36 \pm 0.01$ | $2.7 \pm 0.1$ | $45 \pm 2$ | $48 \pm 2$ | $7 \pm 1$ | (Bakshi et al., 2013; Stracy et al., 2015) |
| DNA polymerase I (PolA) | DNA repair / replication | 128 (monomer) | 500 | $1.05 \pm 0.01$ | $6.6 \pm 0.2$ | $4 \pm 1$ | $61 \pm 2$ | $35 \pm 2$ | (Uphoff et al., 2013) |
| MukBEF (MukB subunit) | chromosome organization/ | 1000 (dimer) | 100 | $0.44 \pm 0.01$ | $1.2 \pm 0.1$ | $58 \pm 2$ | $24 \pm 1$ | $18 \pm 1$ | (Badrinarayana n et al., 2012) |
| DNA Ligase (LigA) | DNA repair / replication | 102 (monomer) | 100 | $1.22 \pm 0.02$ | $6.9 \pm 0.3$ | $7 \pm 1$ | $56 \pm 2$ | $37 \pm 2$ | (Uphoff et al., 2013) |
| Lac repressor (LacI) | gene regulation | 220 (tetramer) | 40 | $0.40 \pm 0.01$ | $3.3 \pm 0.2$ | $41 \pm 3$ | $55 \pm 3$ | $4 \pm 1$ | (Garza de Leon et al., 2017) |
| Heat Unstable protein (HU) | nucleoid associated protein/ gene regulation | 48 (heterodimer) | 30,000 | $0.33 \pm 0.01$ | $12.6 \pm 4.8^\dagger$ | $23 \pm 1$ | $76 \pm 1^\dagger$ | $0.8 \pm 2^\dagger$ | (Gruber, 2014) |
| Histone-like nucleoid structuring protein (H-NS) | nucleoid associated protein/ gene regulation | 87 (dimer) | 20,000 | $0.41 \pm 0.01$ | $8.0 \pm 2.2^\dagger$ | $73 \pm 2$ | $27 \pm 2^\dagger$ | $0.3 \pm 1^\dagger$ | (Katayama et al., 1996) |
| MutS | DNA repair | 248 (dimer) | 100 | $0.37 \pm 0.01$ | $3.7 \pm 1.7^\dagger$ | $26 \pm 2$ | $69 \pm 7^\dagger$ | $5 \pm 1^\dagger$ | (Uphoff et al., 2016) |
| Topoisomerase IV (ParC subunit) | Supercoiling/ decatenation | 366 (heterotetramer) | 80 | $0.40 \pm 0.01$ | $2.7 \pm 1.8^\dagger$ | $37 \pm 2$ | $52 \pm 18^\dagger$ | $11 \pm 7^\dagger$ | (Zawadzki et al., 2015) |
| UvrA | DNA repair | 270<br>(dimer) | 80 | $0.38 \pm 0.02$ | $3.4 \pm 1.7^{\dagger}$ | $43 \pm 2$ | $51 \pm 8^{\dagger}$ | $6 \pm 18^{\dagger}$ | (Stracy et al., 2016) |
| DNA gyrase<br>(GyrA subunit) | Supercoiling | 424<br>(heterotetramer) | 600 | $0.35 \pm 0.01$ | $2.4 \pm 1.8^{\dagger}$ | $55 \pm 1$ | $39 \pm 17^{\dagger}$ | $7 \pm 8^{\dagger}$ | (Stracy et al., 2019) |

<sup>†</sup>Note that in some cases the functional complex contains more than one copy of the subunit which has been labelled, and the complex therefore contains more than one PAmCherry protein. <sup>†</sup>Denotes predicted values based on extrapolation of $D_{free}$ values in shown in Fig. 6.

### Transient DNA-binding events dominate the target search

Comparison of the diffusion coefficient in unperturbed cells versus chromosome-free cells allows quantifying the contribution of non-specific DNA interactions to the observed mobility of DNA-binding proteins during the target search. By simulating molecules rapidly interconverting between freely diffusing (with *D*_*free*_ determined from chromosome-free cells) and DNA-bound (*D*_*bound*_ = 0.04 µm^2^/s) we can establish the fraction of time a protein spends transiently bound to DNA, Φ_transient_binding,_ which best recapitulates the observed mobility of mobile molecules during their target search measured in unperturbed cells (*D*\*_*mobile*_, Fig. 1B). Using this approach we find that Φ_transient_binding_ > 0.5 for RNAP, Pol1, MukB, and LigA, demonstrating that they spend the majority of their search process non-specifically bound to DNA (Φ_transient_binding_ RNAP: 87%, LacI: 93%, Pol1: 61%, MukB: 58%; LigA: 60%) (Fig. 6C). The value for LacI is in good agreement with a previous estimate (Elf et al., 2007). Including the populations of long-lived immobile molecules measured in unperturbed cells (representing molecules likely to be bound at specific DNA target sites, Fig. 1) in the calculations, the total percentage of DNA-bound molecules at any time is even higher (RNAP: 93%, LacI: 96%, Pol1: 63%, MukB: 82%; LigA: 63%) (Fig. 6D). Based on these results, we report a quantitative partitioning of DNA-binding proteins into three distinct states of mobility: long-lived specific binding at DNA target sites, transient non-specific DNA-binding, and free diffusion between DNA strands (Fig. 6C-D).

Using the estimate of α = -0.75 to extrapolate *D*_*free*_ and the measured % of long-lived DNA binding and the measured *D*\*_*mobile*_ values, we performed the same partitioning of diffusive states for all the DNA-binding proteins considered in this study (Table 1). In all cases, the fraction of the target search spent non-specifically bound to DNA was >50%, and for the small nucleoid associated protein HU this estimated fraction was as high as 99%.

## DISCUSSION

Our study demonstrates the ubiquity of transient non-specific DNA interactions for diverse DNA-binding proteins *in vivo*. Despite their different sizes, DNA targets, mobility, and copy numbers in the cell, the target search of all the DNA-binding proteins examined here is dominated by transient non-specific DNA binding. Considering such widespread and frequent non-specific DNA interactions of all types of DNA-binding proteins, an important question is how these in turn affect DNA transactions. Our catalogue of the intracellular mobility of different types of DNA-binding proteins can serve as a general reference and should aid ongoing efforts to generate physical models of the intracellular environment.

Our analysis shows that the chromosome DNA mesh does not constitute a physical barrier for the intracellular motion of proteins (at least up to a protein weight of 100 kDa). In fact, mobile DNA-binding proteins (even large complexes such as RNAP) are enriched in the densest regions of the nucleoid by frequent non-specific DNA interactions. These results demonstrate that the apparent mobility of DNA-binding proteins depends on DNA-binding activity rather than molecular weight, as concluded from previous FRAP experiments (Kumar et al., 2010). While we have found no evidence of a nucleoid sieving effect for DNA-binding proteins during their target search, previous reports have established that large macromolecular complexes which do not bind DNA, such as protein aggregates, 70S ribosomes, and MS2-RNA systems are excluded from the nucleoid (Landgraf et al., 2012; Lindner et al., 2008; Stracy et al., 2015; Stylianidou et al., 2014). Smaller non-DNA-binding complexes such as individual ribosomal subunits appear to diffuse freely through the nucleoid (Sanamrad et al., 2014). Together, these findings are consistent with the view that DNA-interacting proteins can diffuse freely in the whole cell compartment and are enriched within the nucleoid volume due to frequent non-specific interactions with the DNA. Protein hopping or sliding along the DNA can enhance the search efficiency for any individual protein, while overcrowding the chromosome with non-specifically bound proteins would globally reduce the search kinetics due to the obstruction of target sites and sliding collisions (Li et al., 2009). This trade-off likely influenced the evolution of protein abundances and their non-specific DNA binding affinities.

Given the diversity of the proteins we have tested, their target-specific DNA interactions are likely to be very different from each other, suggesting that a more universal interaction plays the largest role in the abundant transient DNA binding during the target search. We speculate that the electrostatic interaction between positively charged functional groups on the surface of the proteins and the largely invariant negatively charged phosphate backbone of the DNA may drive this phenomenon (Kalodimos et al., 2004; Redding and Greene, 2013). Indeed, the surface charge of proteins strongly affects their mobility in cells (Elowitz et al., 1999; Schavemaker et al., 2017), and high intracellular salt concentrations can disrupt DNA-binding *in vivo (Cagliero and Jin, 2013)*. The abundance of non-specific binding also suggests that the percentage of the chromosome occupied by proteins may be higher than expected. Based on the combined percentage of DNA-bound proteins (both specific and non-specific) in Table 1, together with literature estimates of their copy number and DNA footprint, we estimate that at any given time 28% of the chromosome is occupied by the 11 proteins studied (12% long-lived binding and 16% transient binding, see Methods). These proteins represent just a fraction of all DNA-binding proteins, suggesting the total DNA occupancy of the entire proteome is substantially higher. This high occupancy, or chromosome-crowding, highlights the importance of studying protein-DNA interaction in the native cellular environment. Besides the target search, non-specific DNA binding likely also influences the dissociation of proteins from their specific target sites. Several studies have shown that competition with proteins in solution accelerates DNA unbinding due to invasion of a partially-dissociated state (Chen et al., 2015; Gibb et al., 2014; Graham et al., 2011; Loparo et al., 2011). Although this has been demonstrated for exchanges between identical proteins in solution and on DNA, the overwhelming abundance of other DNA-binding proteins and their frequent transient associations with DNA likely contribute significantly to the turnover of DNA-bound proteins in vivo. Thus, non-specific DNA interactions play a crucial role in both the search and the dissociation of DNA-binding proteins.

Beyond these fundamental implications, our chromosome-degradation system has broader potential applications in synthetic biology and has benefits compared to alternative approaches such as chromosome-free minicells. Minicells can be generated by forcing aberrant cell divisions close to the cell poles, however these cells have a perturbed makeup of proteins and contain few DNA binding proteins (Shepherd et al., 2001). In contrast, our chromosome-degraded cells retain the DNA binding proteins and maintain the same cell size and geometry. Moreover, we have shown that DNA-free *E. coli* cells maintain ATP levels and continue to produce plasmid-encoded proteins for several hours, enabling targeted expression of exogenous genes without interference from chromosomal gene expression. Removing all endogenous gene circuitry from *E. coli* cells but maintaining the transcription machinery provides customizable non-viable containers for a range of applications, including expression of synthetic gene circuits, biosensing, and drug delivery (Caliando and Voigt, 2015; Fan et al., 2020; MacDiarmid et al., 2007; Rampley et al., 2017).

## Methods

### Bacterial strains, plasmids and growth

Bacterial strains and plasmids are listed in Table S1. All experiments were performed in *E. coli* TB28 background strain (MG1655 *ΔlacIZYA*) (Bernhardt and de Boer, 2005). PAmCherry fusion proteins expressed from their endogenous chromosome loci were previously characterized: RNAP, HU and HN-S (Stracy et al., 2015), LacI (Garza de Leon et al., 2017), Pol1 and LigA (Uphoff et al., 2013), UvrA (Stracy et al., 2016), MutS (Uphoff et al., 2016), ParC (Zawadzki et al., 2015), MukB (Badrinarayanan et al., 2012), GyrA (Stracy et al., 2019). Fusions were moved to *E. coli* TB28 strain by P1 transduction. Construction of plasmids expressing LacI-PAmCherry or LacI mutant are described in (Garza de Leon et al., 2017). Unconjugated PAmCherry was produced from the plasmid pBAD\HisB PAmCherry1 (Endesfelder et al., 2013). ParB-mCherry was produced from pSN70 plasmid (Nolivos et al., 2019). The *I-SceI* cut site (*I-SceI*^*CS*^) is followed by *cat* gene (chloramphenicol resistance) flanked by *frt* sites as described previously (Lesterlin et al., 2013). *I-SceI*^*CS*^ was inserted in two chromosome loci by λ-Red recombination (Datsenko and Wanner, 2000); *ilvA* (3953 kb) close to the origin of replication, and *ydeO* (1580 kb) in the *terminus* region. Using sequential P1 transduction, we constructed the OT strain (for *Ori-Ter*) carrying *ilvA::I-SceI*^*CS*^ and *ydeO::I-SceI*^*CS*^. After each transduction round, the *cat* gene was removed using Pcp20 plasmid (Datsenko and Wanner, 2000). P1 transduction was also used to transfer *recA-* mutation *recAT233C-Tet* or *minC-Ypet* allele (Bisicchia et al., 2013) alleles. Unless otherwise stated, cells were grown at 30°C in M9 medium supplemented with glucose (0.2%). When appropriate, growth media were supplemented with Ampicillin (Ap) 100 µg/ml, Chloramphenicol (Cm) 20 µg/ml or Kanamycin (Kn) 50 µg/ml.

### Sample preparation for microscopy

OT strains carrying two *I-SceI*^*CS*^ were transformed with pSN1 plasmid carrying the *I-SceI* gene under the control of the *P*_*lac*_ promoter and plated on LB agarose plates containing 0.2 % glucose and ampicillin at 30°C. Transformant clones were propagated on LB agarose plates containing 0.2 % glucose and ampicillin. Transformation was performed *de novo* before each experiment since strains carrying *I-SceI*^*CS*^ and the pSN1 plasmid exhibit genetic instability due to leaky *I-SceI* expression causing unrepairable DNA double-stranded breaks in the *recA-* strain. For each strain, a single colony was inoculated in M9 minimal medium supplemented with 0.2% glucose and ampicillin and incubated overnight at 30°C with agitation (140 rpm). The next day, overnight cultures were diluted and grown to early exponential phase (OD_600nm_ ∼ 0.2). 0.2% arabinose was added to induce the production of I-SceI endonuclease and initiate chromosome degradation in the *recA-* strains. Cultures were incubated at 30°C with agitation for the duration indicated in the text and figures (120 min for complete DNA degradation) before microscopy. For control experiments in fixed cells, 2.5% paraformaldehyde was added to the growth media for 1 hour prior to imaging. Cell filamentation was induced by addition of cephalexin at final concentration of 5 µg/ml.

The cell suspension was concentrated by centrifugation (benchtop centrifuge at 6000 rpm), removal of the supernatant and resuspension in 1/10^th^ of the initial sample volume. Cells were immobilized on pads of 1% low-fluorescence agarose (Biorad) in M9 medium with 0.2% glucose as previously described (Lesterlin and Duabrry, 2016). For PALM microscopy 0.17 mm thickness coverslips were heated in an oven to 500°C to remove any background fluorescent particles before use. For quantification of chromosome degradation and MinC-Ypet oscillation by wide-field epifluorescence imaging, DNA staining was performed by incubating the cell suspension for 15 min with 2 4’,6-diamidino-2-phenylindole (DAPI) at 4 µg/ml prior to cell concentration and imaging. For multi-color imaging of the nucleoid and PAmCherry fusions, we stained DNA with 500 nM SytoGreen for 15 min before imaging (because DAPI excitation would cause photoactivation of PAmCherry).

### Wide-field epifluorescence microscopy imaging

Wide-field epifluorescence microscopy imaging of DAPI-stained cells was carried out on an Eclipse Ti-E microscope (Nikon), equipped with 100s/1.45 oil Plan Apo Lambda phase objective, Flash4 V2 CMOS camera (Hamamatsu), and using NIS Elements software for image acquisition. Acquisition was performed in phase contrast and epifluorescence mode using 50% power of a Fluo LED Spectra X light source at 405 nm and 560 nm excitation wavelengths for DAPI and ParB-mCherry, respectively. Wide-field imaging of MinC-Ypet was carried out on a Nikon Eclipse TE2000-U microscope equipped with a 100X objective, CCD camera (Cool-SNAP by Photometrics) and Metamorph 6.2 acquisition software. Time-lapse movies were acquired in phase contrast and epifluorescence at 2-s intervals with 50 ms exposure for MinC-Ypet at 30°C.

#### Widefield epifluorescence image analysis

Cells were automatically detected using the MicrobeJ plugin for Fiji (Ducret et al., 2016). Intracellular DAPI or ParB-mCherry mean fluorescence intensity (a.u.) was automatically extracted and plotted using the MicrobeJ results interface. For analysis of MinC oscillation, cells were outlined using the MATLAB-based tool MicrobeTracker (Sliusarenko et al., 2011). The fluorescence signal was integrated across the cross-section of each cell to generate a one-dimensional fluorescence profile in each frame. The fluorescence signal was normalized to the total fluorescence in each frame to remove photobleaching effects and facilitate MinC-Ypet localization analysis. The fluorescence signals obtained from each cell were further analyzed by generating kymographs using custom MATLAB code. The width of the kymograph corresponds to the cell length L. We integrated the fluorescence intensity for both cell halves at in each frame,

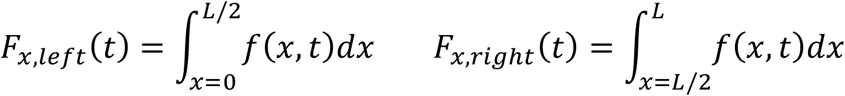

By fitting the data to a trigonometric function the oscillation period is calculated from the angular frequency 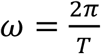

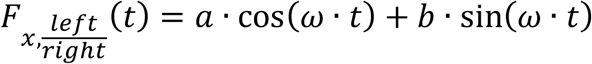

The time-averaged concentration profile of MinC is obtained by integration of the entire kymograph over all frames,

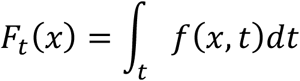

For analyzing MinC oscillations in filamentous cells, a slightly modified kymograph analysis was used. The MinC concentration profile was determined as described above. The positions of the fluorescence minima *xmin* were used to split the kymographs into several stripes. The overall oscillation period *Tm* was calculated as the average of all oscillation periods determined for each stripe in the kymograph. The oscillation wavelength was determined from the distance between two neighboring peaks. Depending on the length of the cell and the number of oscillations a set of wavelengths was determined from which a mean wavelength was calculated as

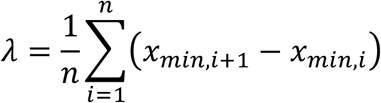

Where the number *n* of oscillations corresponds to the number of peaks – 1.

### Live-cell photoactivated single-molecule tracking

Live cell photoactivated single-molecule tracking was performed on a custom-built total internal reflection fluorescence (TIRF) microscope built around the Rapid Automated Modular Microscope (RAMM) System (ASI Imaging) as previously described (Uphoff, 2016). PAmCherry activation was controlled by a 405 nm laser and excited with 561 nm. All lasers were provided by a multi-laser engine (iChrome MLE, Toptica). At the fiber output, the laser beams were collimated and focused (100x oil immersion objective, NA 1.4, Olympus) onto the sample under an angle allowing for highly inclined thin illumination (Tokunaga et al., 2008). Fluorescence emission was filtered by a dichroic mirror and filter (ZT405/488/561rpc & ZET405/488/561NF, Chroma). PAmCherry emission was projected onto an EMCCD camera (iXon Ultra, 512×512 pixels, Andor). The pixel size was 96 nm. Transmission illumination was provided by an LED source and condenser (ASI Imaging and Olympus). Sample position and focus were controlled with a motorized piezo stage, a z-motor objective mount, and autofocus system (MS-2000, PZ-2000FT, CRISP, ASI Imaging). Movies were acquired under continuous laser excitation with exposure times of 15 ms or 5 ms for 20,000 frames at 20°C. Camera readout was 0.48 ms giving frame intervals of 15.48 ms or 5.48 ms, respectively. We also recorded a transmitted light snapshot for segmenting cells in each movie. For imaging SytoGreen, snapshots with 488 nm excitation with a 50 ms exposure time were acquired prior to PAmCherry imaging.

#### Localization and tracking

Single-molecule-tracking analysis was performed using custom-written MATLAB software (MathWorks) as previously described (Uphoff et al., 2014): fluorophore images were identified for localization by band-pass filtering and applying an intensity threshold to each frame of the movie. Candidate positions were used as initial guesses in a two-dimensional elliptical Gaussian fit for high-precision localization. Free fit parameters were x-position, y-position, x-width, y-width, elliptical rotation angle, intensity, background. Localizations were segmented based on cell outlines obtained from MicrobeTracker applied to the brightfield snapshots. Single-particle tracking analysis was performed by adapting the MATLAB implementation of the algorithm described in (Crocker and Grier, 1996). Positions were linked to a track if they appeared in consecutive frames within a window of 5 pixels (0.48 µm). When multiple localizations fell within the tracking window, tracks were linked such that the sum of step distances was minimized. We used a ‘memory’ parameter of 1 frame to allow for transient disappearance of the fluorophore within a track due to blinking or missed localization.

#### Measuring the diffusion of tracked molecules

We determined the mobility of each molecule by calculating an individual apparent diffusion coefficient, 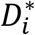, from the one-step mean-squared displacement (MSD) of the track using:

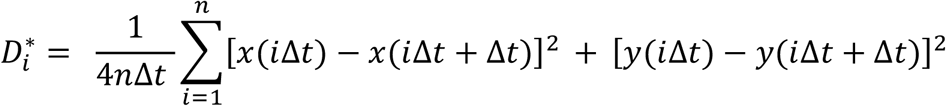

Where *x*(*t*) and *y*(*t*) are the coordinates of the molecule at time *t*, the frame time of the camera is *Δt*, and *n* is the number of frames over which the molecule is tracked. For a molecule diffusing with an apparent diffusion coefficient *D*\*, the probability of measuring a 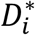 by tracking it over *n* frames, is given by (Vrljic et al., 2002):

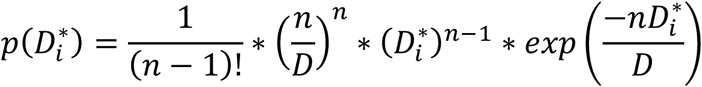

In order to determine the apparent diffusion coefficient, *D*\*, from the population of individual single-molecule 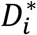 values, longer tracks were truncated after 5^th^ localization (i.e. *n* = 4). The 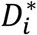 distribution was then fitted with the equation for *n* = 4 :

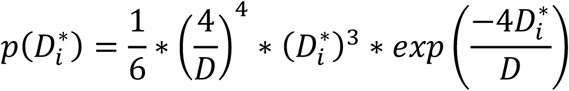

Fits were performed using maximum likelihood estimation in MATLAB. For unperturbed cells the protein diffusion distributions were fit with a model containing two molecular species with diffusion coefficients 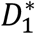 and 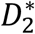: representing immobile molecules bound to DNA for the entire trajectory, and mobile molecules diffusing and binding only transiently to DNA:

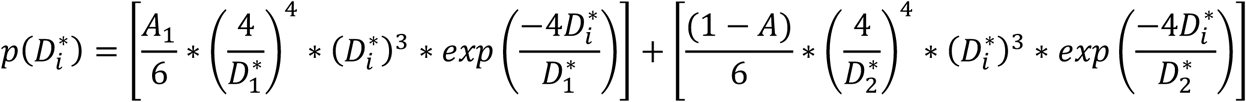

where *A*and 1 − *A* are the fraction of molecules found in each state. The localization uncertainty, σ_loc_, manifests itself as a positive offset of σ_loc_^2^/Δt in the *D*\* value (Michalet and Berglund, 2012). Based on the estimated localization uncertainty of ∼35 nm for our measurements, we expected a positive shift in the mean *D*\* value of immobile molecules to ∼0.7 µm^2^s^-1^. Where indicated error bars represent 95% confidence intervals obtained from fitting the D* distribution for 1000 bootstrap resamplings with replacement of individual segmented cells. For each bootstrap the tracks within the sampled cells were pooled and fitted as described above.

To plot maps of tracks from mobile and immobile molecules, we used a threshold of 0.15 µm^2^s^-1^ to separate the populations.

#### Monte Carlo diffusion simulations

The apparent diffusion coefficients determined experimentally through particle tracking do not take into account three-dimensional confinement in the bacterial cell. We followed a similar rationale as before to remove this bias (Uphoff et al., 2013): we simulated Brownian motion confined within 3D cell volumes obtained from the segmented 2D brightfield images. The distance from the midline to the cell edge was used as the radius of a cylindrical volume for each length segment of a cell. For each cell diffusion simulations of the same number of molecules as measured experimentally were performed. Each 15 ms frame was split into 100 sub-frames with Gaussian-distributed displacements in each sub-frame. Each molecule trajectory was given a random starting time to mimic stochastic photoactivation, and a duration sampled from an exponential distribution with a mean time equal to our experimentally determined photobleaching lifetime (85 ms). The sub-frame distributions were then averaged to give a position for each frame, and a localization error sampled from a Gaussian distribution with σ_loc_ = 35 nm was added. The list of simulated localizations, with their corresponding frame numbers was then analyzed using the same algorithms with the same settings as for experimental data. The best estimate for the unbiased diffusion coefficient was determined by running the simulations for different *D* values between 0 and 10 μm^2^s^-1^ and selecting inputted *D* value from the simulated *D*\* distribution which best approximates (based on the least squares error) the experimentally obtained *D*\* distribution. Since diffusion coefficients in DNA-free cells were much higher than in unperturbed cells we also performed experiments and simulations for Pol1 and MukB at 5.48 ms exposure times to verify the same underlying unbiased diffusion coefficients were obtained independent of the data acquisition conditions. 95% confidence intervals were estimated by fitting the experimental *D*\* distribution for 1000 bootstrap resamplings with replacement of individual segmented cells as described previously. Simulations were then performed to determine inputted D value which best approximates the higher and lower confidence bounds from the experimentally determined *D*\* values.

We hypothesized that the observed diffusion of DNA-binding proteins in unperturbed cells represented mobile molecules interconverting between D_free_ and D_bound_ states. By comparing diffusion in unperturbed and DNA-free cells, it is possible to estimate the relative occupation of the states but not the absolute duration a molecule spends in each state. To simulate molecules interconverting between these states, we used the D_free_ value based on the simulations of DNA-free cells, and a D_bound_ value of 0.04 μm^2^s^-1^. Because the D_mobile_ population appears as a single species the interconversions must occur on a timescale below the observation window per track (75 ms, 5 frames of 15 ms). We therefore simulated the duration of D_bound_, t_bound_, by randomly sampling from an exponential distribution with a mean of 1 ms. We performed simulations for a range of ratios of durations in the D_free_ and D_bound_ states by varying, t_free_, the duration of free diffusion between binding events. Using least squares optimization, we determined the ratio which best recapitulated the experimental *D*\*_*mobile*_ value determined from fitting the experimental D* distribution.

#### Chromosome occupancy calculations

To estimate the percentage of the chromosome occupied by proteins we used literature estimates of the DNA footprint of each protein. RNAP (70bp, (Ross and Gourse, 2005); HU (36bp, Gruber, 2014); H-NS (30bp; van der Valk et al., 2017); DNA gyrase (100bp, Reece and Maxwell, 1991). Where no DNA footprint estimates could be found we assumed a footprint of 10bp. The total bp occupied was calculated by the molecules/cell multiplied by the total fraction binding (including stable binding and transient binding) in Table 1, and the DNA footprint, giving 1.96Mb of DNA. Under the minimal media growth conditions in this study there are on average 1.5 chromosomes per cell, totaling 6.9Mb of DNA (Wang et al., 2005).

## ACKNOWLEDGMENTS

The authors thank Sophie Nolivos for providing pSN1 plasmid;

## Author contributions

M.S., S.U. and C.L. conceived the project, designed the study and wrote the paper with equal contributions; M.S., S.U, J.S. and C.L. performed the experiments and analyzed the data. M.S., S.U., DJ.S., A.K. and C.L. provided funding and gave intellectual input throughout the project.

## Data and materials availability

Raw data is available from Mendeley Data. Materials are available from the corresponding authors upon request.

## SUPPLEMENTARY FIGURES

**Figure S1:**
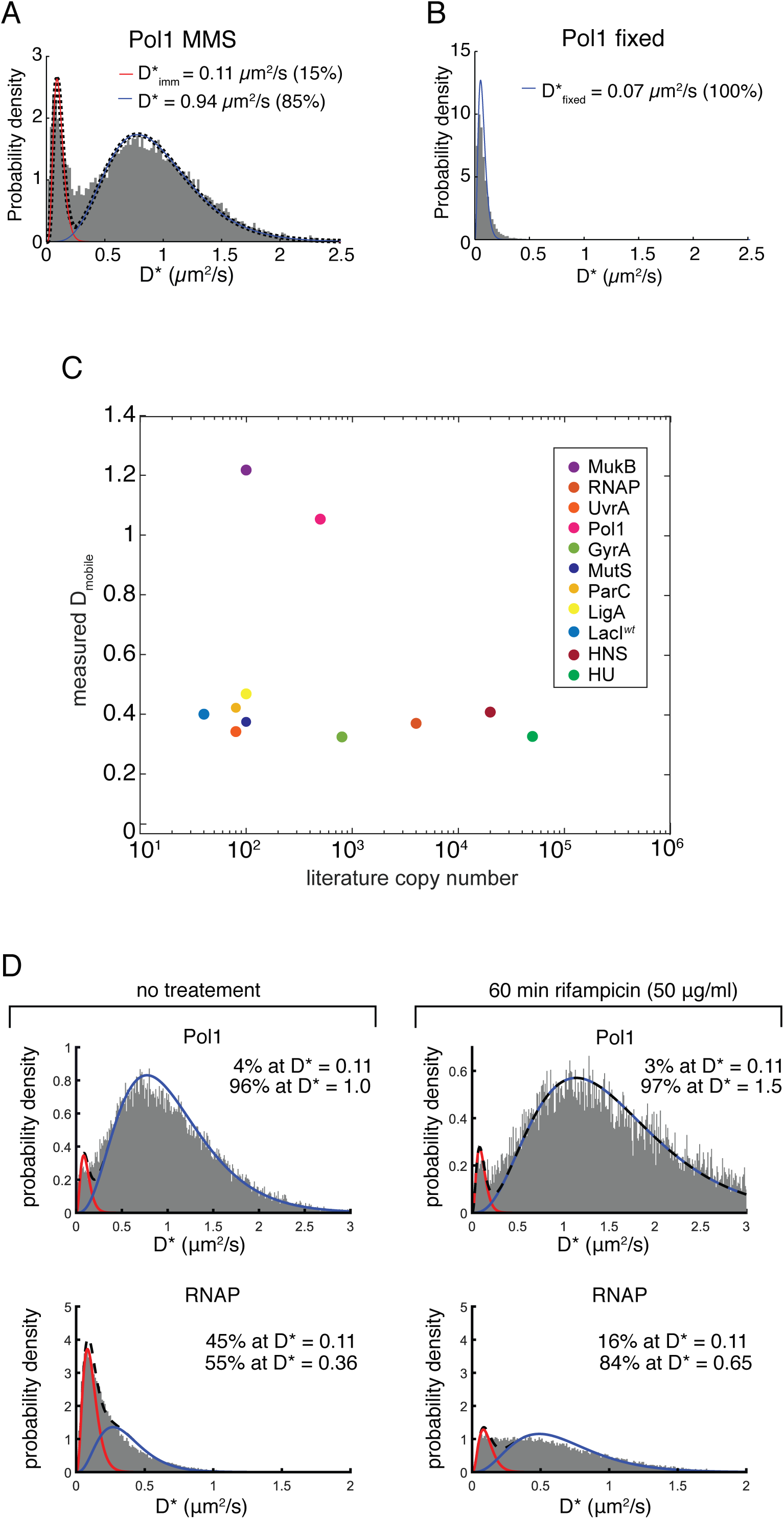
(A) D* histograms and model fit of PolA-PAmCherry in cells treated with methyl methanesulfonate (MMS). (B) D* histograms of PolA-PAmCherry in fixed cells, fitted with a model of immobile molecules only. (C) Scatter plot of experimentally determined D_mobile_ diffusion of 12 DNA-binding proteins against their intracellular copy number. The fitted D_mobile_ value extracted from a 2 species fit to the histograms of apparent diffusion coefficients, D*, presented in Figure 1. The copy number estimates are from literature sources are presented in Table 1. (D) Distributions histograms of apparent diffusion, D*, of three DNA-binding proteins: DNA polymerase 1 (Pol1) and RNA polymerase (RNAP) before (left) and after 60 mins treatment with 50 µg/ml rifampicin (right). Distributions are fitted with a 2-species model of immobile (in red) and mobile molecules (in blue).

**Figure S2:**
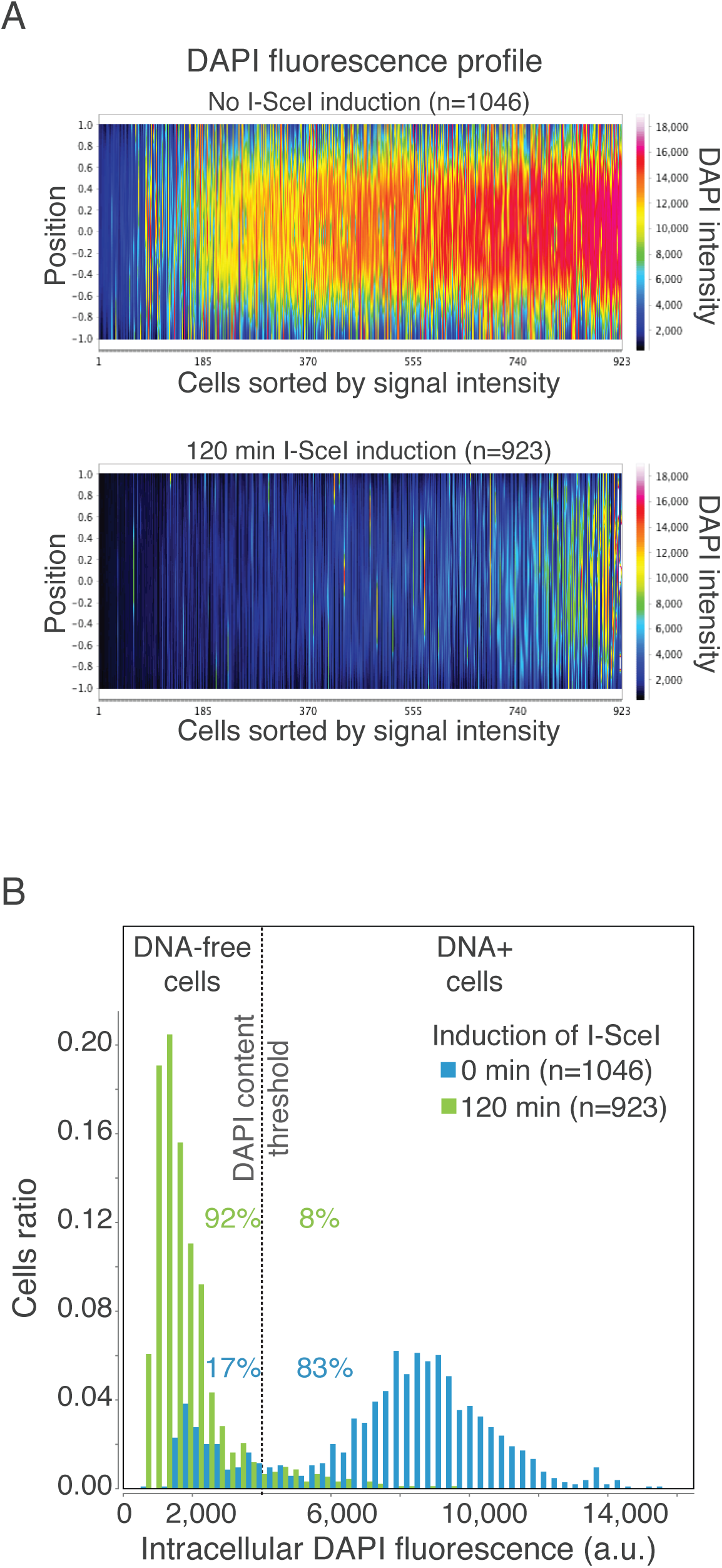
Quantification of DNA degradation efficiency. (A) Fluorescence profiles show the distribution and intensity of DAPI signal in individual cells normalized by the cell length and sorted from left to right by increasing mean intensity. Fluorescence profile of OT / pSN1(p*P*_*BAD*_-*I-SceI*) strain before (Top panel) and 120 min after I-SceI induction (lower panel) are shown. (B) Histograms of DAPI-stained DNA intracellular fluorescence in OT / pSN1(PBADI-SceI) cell population, before (blue bars) and 120 min after I-SceI induction (green bars). The percentages of cells below and above a DAPI content threshold (grey dash line) are shown. Before I-SceI induction, 17 % of cells already exhibit DNA loss likely due to the leakiness I-SceI expression from PBAD promoter. 120 min after arabinose-induced I-SceI expression, this proportion increases to 92 % of the population, with 8 % of cells still containing DNA (n= cells analyzed).

**Figure S3:**
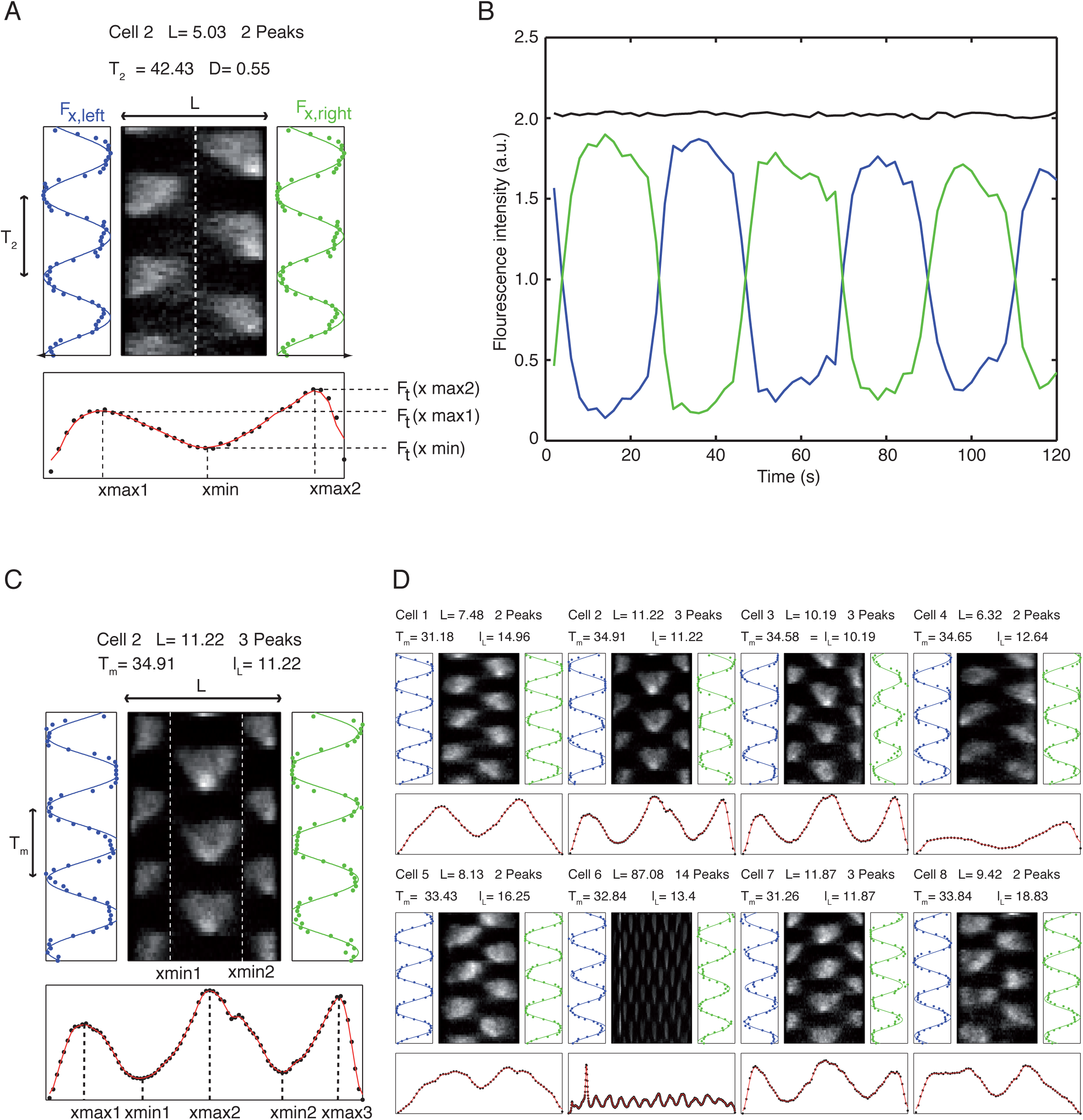
(A) Kymograph and concentration profiles of the fluorescence signal of MinC protein in exponentially growing *E. coli* cells. The width of the kymograph corresponds to the length L of the cell. Upon vertical splitting of the kymograph (white dashed line) and integration over space (x), the time-dependent intensity signals Fx,left(t) and Fx,right(t) are obtained. The oscillation period T2 can be calculated by periodic fitting (blue and green lines on the left and right of the kymograph). Upon integration over time (t), a spatial concentration profile of the MinC proteins is obtained (black data points below the kymograph) and fitted (red curve). The depth D of the profile is calculated from the heights of the maxima and the minimum. (B) Oscillation of fluorescence over time in cell halves. When the time-dependent fluorescence is integrated, an almost perfect constant line is obtained (black curve). This is due to the normalization of the intensity of each pixel in respect to the total fluorescence signal at each time point. In consequence, the two periodic curves for the left and right kymograph halves are perfectly symmetric (blue and green curve). (C) Kymograph and concentration profiles of the fluorescence signal of MinC protein in filamentous cells. In filamentous cells, the time-averaged concentration profile can show more than two peaks. The distance between two peaks corresponds to half an intrinsic wavelength of the Min system. For the calculation of the oscillation period Tm in filamentous cells, the kymograph is split along the position of the minima xmin (white dashed lines in the kymograph) resulting in several stripes with a periodic pattern. Integration along x provides a periodic functions Fx,i(t) from which oscillation period Ti can be calculated. Tm represents the average for these multiple oscillation periods. (D) Kymographs of different filamentous cells show different number of oscillations. Most of them show only a single (cells 1, 4, 5 and 8) or double oscillation (cells 2, 3 and 7). In the given example one cells shows 14 peaks, which corresponds to 13 oscillations and thus 6.5 wavelengths. The respective wavelength can be then calculated from the length of the cell and the number of oscillations.

**Figure S4:**
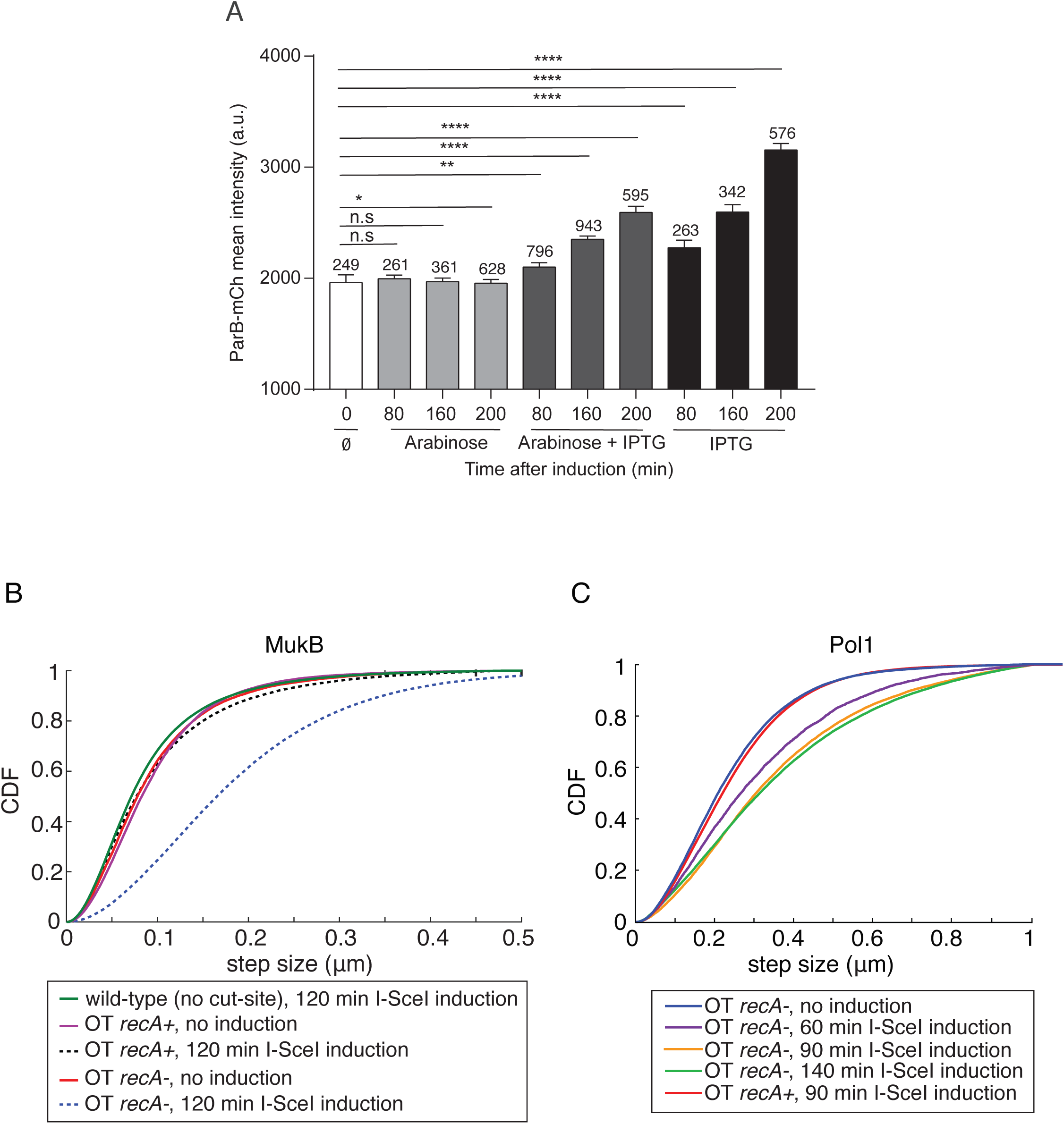
(A) Maintenance of protein synthesis activity upon induction of DNA degradation. Quantification of ParB-mCherry intracellular signal (a.u., arbitrary unit) produced from IPTG inducible pSN70 plasmid (*P*_*lac*_ *I-SceI*) in OT strain. I-SceI expression from pSN1 plasmid is induced by arabinose 0.2 %. Chromosome degradation alone has little impact on ParB-mCherry production. Over the course of the experiment (200 minutes), ParB-mCherry production is induced by IPTG with or without chromosome degradation. Error bars indicate the standard error and n = the numbers of cells analyzed. Two tailed P-values from Mann-Whitney non-parametric test are indicated by (n.s) non-significant P-value > 0.05, * for P-value < 0.05, ** for P-value < 0.01 and **** for P-value < 0.001. (B) Cumulative distribution plot of tracked MukB-PAmCherry trajectory step size in cells before and 120 mins after I-Scel induction. Protein diffusion remains unchanged *recA* proficient cells (*recA+*) before (magenta line) after induction (black dashed line), whereas diffusion increases after induction in *recA-* cells (blue dashed line). (C) Cumulative distribution plot of tracked Pol1-PAmCherry trajectory step size in (recA-) cells before and after I-Scel induction. Protein diffusion increases with I-Scel induction time from 0 to 90 mins, but increases only modestly beyond 90 mins induction.

**Figure S5:**
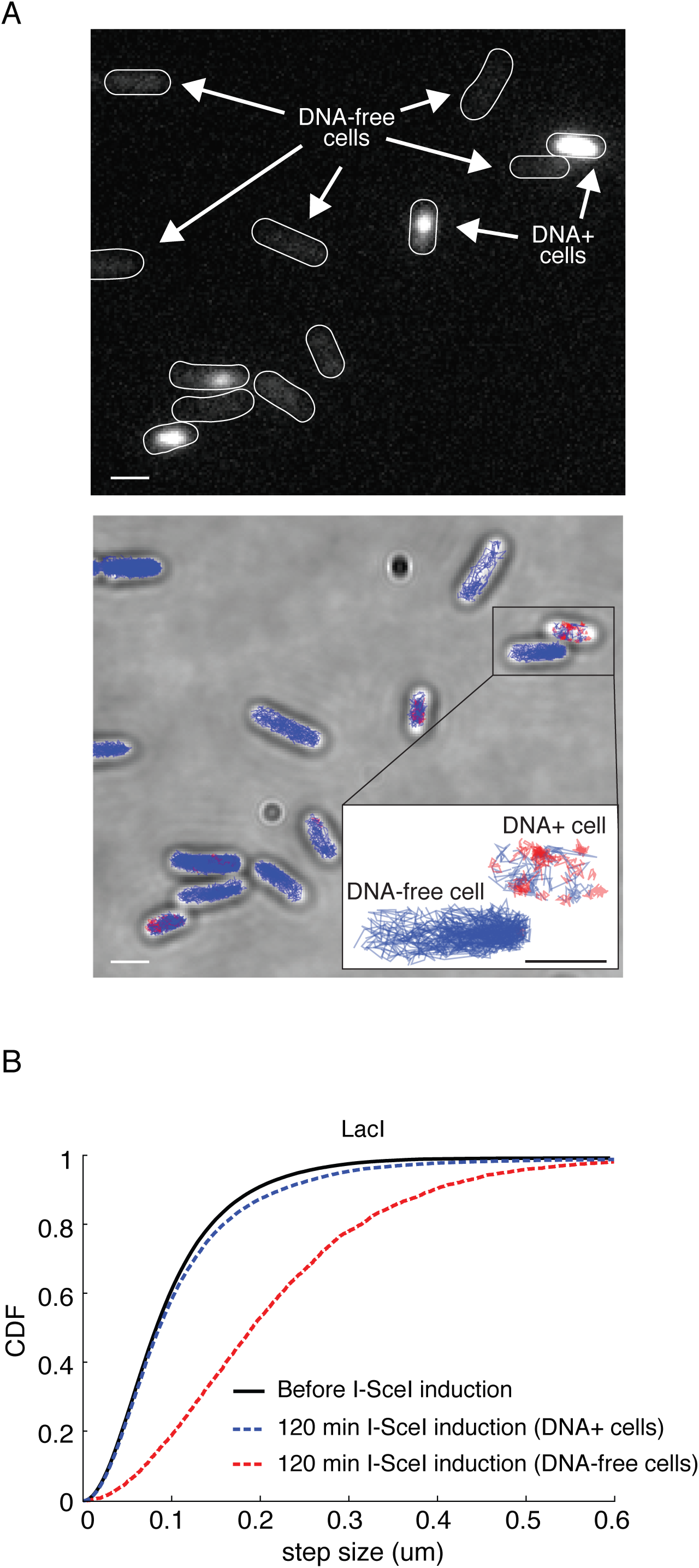
A fraction of cells did not undergo full DNA degradation and these cells showed little change in the diffusion profiles. (A) Fluorescence image of SytoGreen stained DNA in cells after 120 mins of I-SceI induction showing DNA+ and DNA-free cells (top). The brightfield image of the same cells overlaid with the categorized trajectories of RNAP-PAmCherry tracks with immobile molecules in red and mobile molecules in blue (bottom). The lower-right insert presents a zoom of one DNA+ and one DNA-free cell. (B) Cumulative distribution of LacI-PAmCherry trajectory displacement steps in cells having high (DNA+ cells in blue) or low (DNA-free cells in red) SytoGreen fluorescence compared to the unperturbed cells (in black).

**Figure S6:**
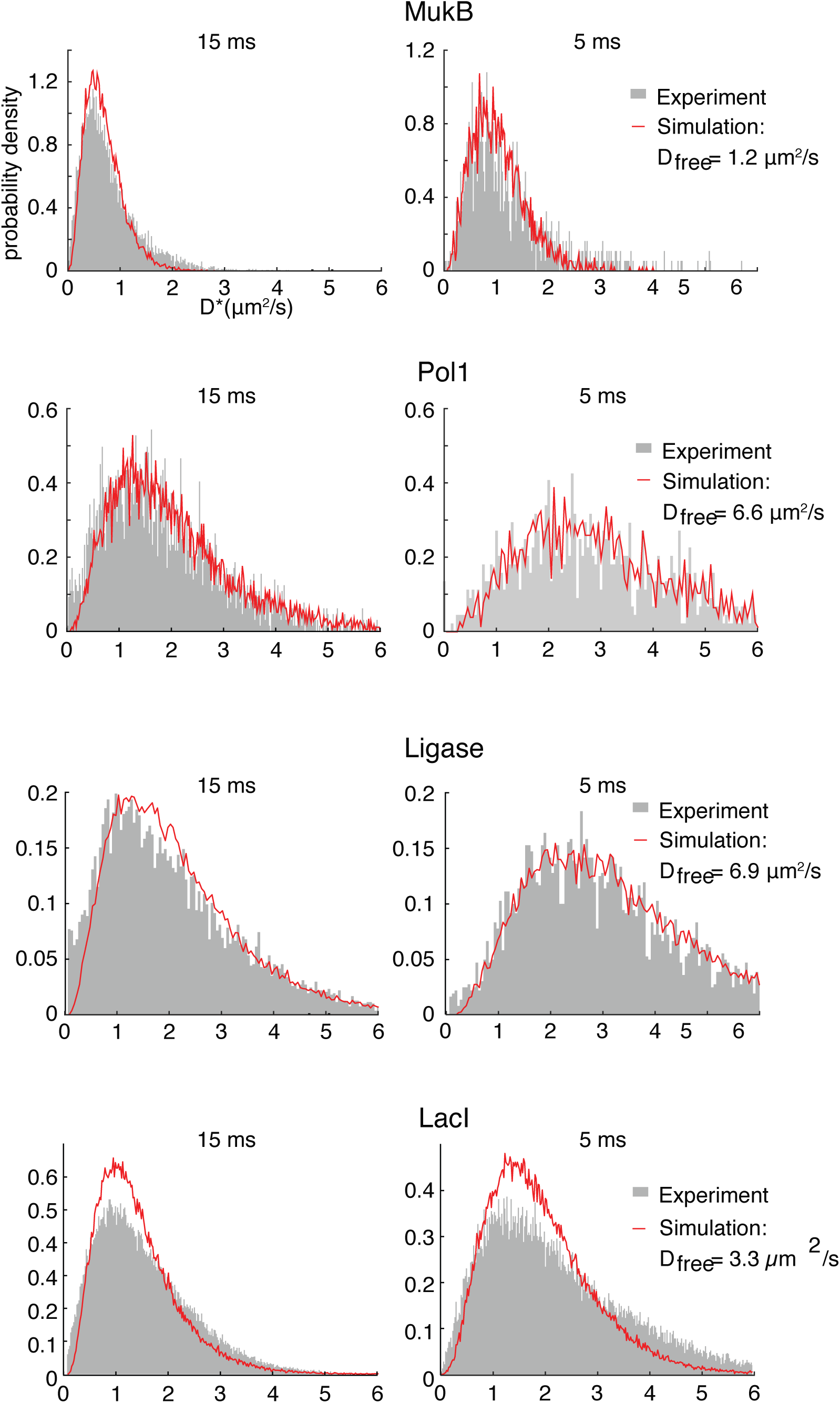
D* distribution (grey bars) of MukB-PAmCherry, Pol1-PAmCherry, LigA-PAmCherry and LacI-PAmCherry in DNA-free cells 120 min after I-SceI induction measured with an exposure time of 15 ms (left) and 5 ms (right). The D* distribution generated from simulated molecule trajectories with the same diffusion coefficient, D_free_, at 15 ms (left) and 5 ms (right) exposure times.

**Table S1.**
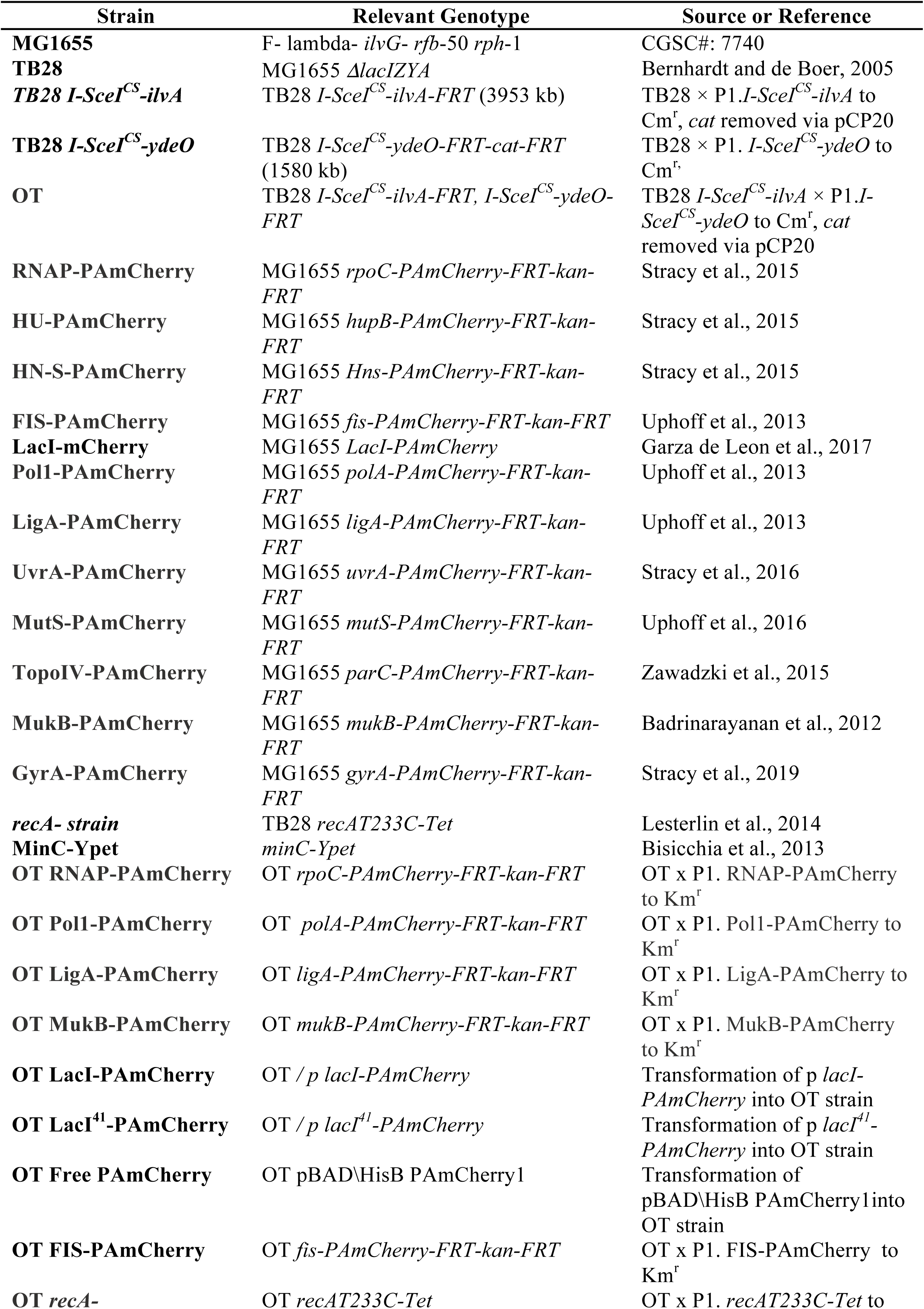
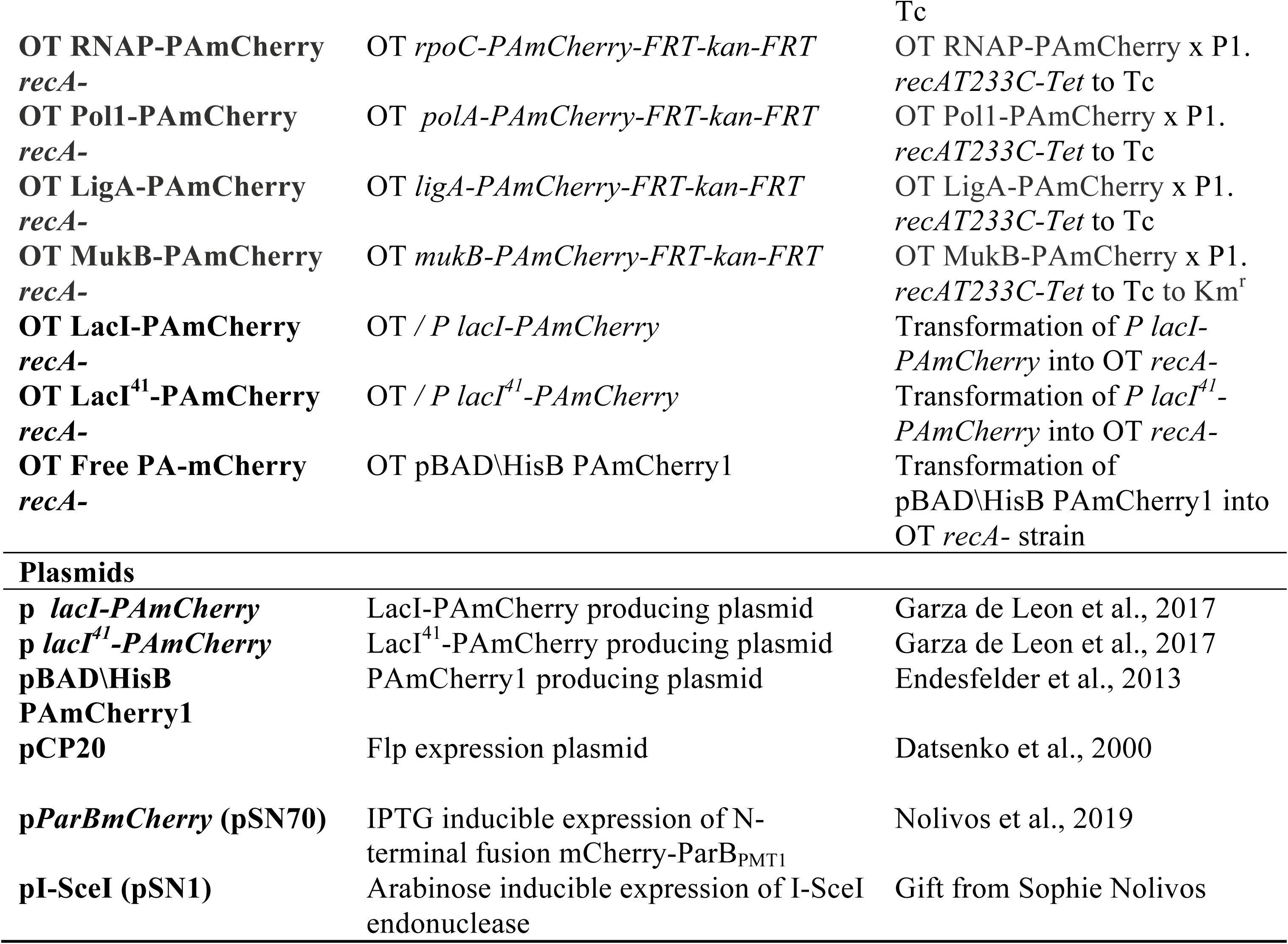
Strains and plasmids used in this study.

